# The benefits of merging passive and active tracking approaches: new insights into riverine migration by salmonid smolts

**DOI:** 10.1101/2021.07.26.453807

**Authors:** Louise Chavarie, Hannele M. Honkanen, Matthew Newton, Jessie M. Lilly, Hannah R. Greetham, Colin E. Adams

## Abstract

The process of smolting is a critical phase in the life-cycle of anadromous salmonids and it has been associated with substantial rates of mortality. Survival during freshwater and marine migration is known to have population level effects, thus an understanding of the patterns of mortality has the potential to yield important insights into population bottlenecks. Despite important advancements in tracking techniques, the specifics of mortality events in anadromous salmonids during their initial migration to sea remains somewhat elusive. Here, we develop a framework combining spatial and temporal detections of smolt riverine migration from two tracking techniques, which enable inferences to be made about mortality locations, causes, and rates. In this study, we demonstrate that during their initial riverine transitional phase, smolts were particularly vulnerable to predators. Specifically, avian predation appeared to be the main cause of mortality (42%), although piscine predation events were not trivial (14%). Our results suggested some direct and indirect tagging-induced mortality (e.g., through increased predation vulnerability), which highlights the importance of determining tagging mortality in a telemetry study to ensure adequate interpretation of migration success. Overall, by estimating migration loss and its variability, our study framework should help to guide management actions to mitigate the widespread population declines these species are currently facing.

## Introduction

Atlantic salmon (*Salmo salar*) and sea trout (the anadromous form of the brown trout complex; *Salmo trutta*) are salmonid species of biological, cultural, and economic importance, but these species have experienced dramatic declines in recent decades (ICES 2018, ICES 2011). Despite various attempts to mitigate anthropogenic stressors and mortality rates, Atlantic salmon and sea trout populations throughout most of geographic range of these fish are currently at or near record low abundances (ICES 2017, 2018). The anadromous life-history tactic of both species results in migration between freshwater and marine habitats, which exposes individuals to multiple threats (e.g., migration barriers, diseases, pollution, overexploitation, parasites, climate change, and aquaculture impacts; Forseth et al. 2017, Parrish et al. 1998). Smolts (the seaward migrating stage of the life cycle) are particularly vulnerable during their first migration to sea because they are increasingly mobile and traverse a high-risk landscape that exposes them to predators (Ward and Hvidsten 2011, Ward et al. 2008). However, the location, timing, and relative proportion of the total mortality attributable to Atlantic salmon and sea trout populations during their initial migration remains elusive (Chaput et al. 2018). This lack of an understanding of migration losses is highly challenging for practical management and for policy development.

Recent advances in the development of electronic tags that transmit information that enables tracking of animals across broad- and fine-spatial and temporal scales have made significant contributions to the understanding of salmonid behaviour and migration (Drenner et al. 2012, Hussey et al. 2015). Our ability to track aquatic organisms often varies with the species studied and the habitat they inhabit. There are trade-offs among operating requirements, transmission quality, and tracking capabilities in the devices used to track animals; these become more complex in species that migrate between freshwater and marine systems (Leander et al. 2019). Currently, electronic tags are the leading technology to track fish with high temporal and spatial resolution in the wild (Hussey, et al. 2015). Radio telemetry has been commonly used in rivers, since it allows the determination of fine-scale movements and mortality (Jepsen et al. 2019, Keefer et al. 2012, Wertheimer and Evans 2005). However, radio telemetry cannot be used to track animals in salt water or deep-water environments, which is a major limitation when studying an anadromous migration (Cooke et al. 2013).

When freshwater and marine migrations by anadromous salmonids are investigated, passive acoustic telemetry offers a good alternative (McMichael et al. 2010). The main issue with passive acoustic telemetry is that tagged individual must move within the detection range of an acoustic receiver to be detected (Both et al. 2012, Bruneel et al. 2020). This issue can be overcome by deploying a sufficient number of receivers to achieve complete coverage of a given area. For example, when the direction of migration is known or constrained in certain areas, using an array of receivers in a grid positioning system in relatively enclosed study areas (e.g., lakes and estuaries) or by using by an array of receivers organised as a gate through which fish are likely to travel, have both been proved to be efficient methods to detect organisms (Heupel et al. 2006, Kraus et al. 2018, Roy et al. 2014). Despite being well suited for marine or lake environments, these solutions to increase the resolution of the data do not necessarily work well in rivers systems. Rivers are often shallow, sinuous and frequently with fast-flowing water that results in a noisy environment that makes tag detection challenging; thus increasing the number of receivers for better detection resolution is not necessarily feasible (Bergé et al. 2012). Combining approaches with different trade-offs (e.g., using PIT tagging or radio telemetry along with acoustic telemetry) may circumvent some of the challenges of tracking in riverine systems, but would require double tagging individuals or running programs in parallel (Dainys et al. 2018, Furey et al. 2016, Jepsen, et al. 2019, Schwinn et al. 2017).

As a result of some of these shortcomings from passive acoustic telemetry associated with rivers, we revisited the concept of active acoustic tracking in association with passive acoustic telemetry. Active acoustic tracking is not a new concept (e.g., Gauld et al. 2016, Halfyard et al. 2012). Active acoustic tracking has been considered to be a detection method with some advantages, it is likely to provide: 1) a high frequency of detections of animal positions, 2) position estimates that are not limited to areas that are in range of fixed receiver stations, and 3) relatively precise animal positions (Brownscombe et al. 2019). However, because active tracking is labour intensive and as it often restricts the sample size compared to a passive tracking approach (e.g., frequently only one animal can be tracked at a time and the duration over which animals can be tracked is usually limited), methods using fixed acoustic receivers have been favoured in telemetry studies (Brownscombe, et al. 2019, Fetterplace et al. 2016). Novel active tracking techniques have been recently developed, such as autonomous robotic technology or receivers attached to large, free ranging animals to enable detection of individuals of the target species (Carlon 2015, Ennasr et al. 2020), none of which are currently suitable for most rivers. Thus, to counter the limitations of existing passive telemetry in small to moderate sized riverine systems and enhance the collected information on smolts during their freshwater migration phase, we developed a novel framework that integrates fixed acoustic receivers and active tracking by systematic canoe transects. The purposes of our study were to: 1) test the relative efficiency of each tracking methodology (i.e., passive and active) separately and together, 2) use the combined active-passive approach to define and quantify the locations, causes, and rates of mortality of salmonid smolts during their riverine migration phase, and 3) determine the fine-scale behaviour related to migration (e.g., time of day travelling and residency).

## Material and methods

### Study Site and Tagging Procedures

The River Endrick (Scotland) is a major inflowing tributary to Loch Lomond, a medium sized lake in the catchment through which fish must migrate to reach the sea (Fig. 1). Wild Atlantic salmon and sea trout smolts (there is no contemporary stocking of these species there) were captured using a 1.2 m diameter rotary screw trap in the middle reaches of the River Endrick, 12.33 km upstream of the river mouth (56 ° 2’ 58’’N; 004 ° 26’ 27’’ W) where it discharges into Loch Lomond. The rotary screw trap was in constant operation (both day and night) over the study period. Between April 13^th^ and 20^th^ 2020, a total of 135 Atlantic salmon and 23 sea trout smolts were tagged with VEMCO V7-2L 69kHz (VEMCO Ltd, Halifax, Canada) tags. To minimize the tag burden, only smolts in excess of 130 mm fork-length were tagged. Smolts were anaesthetised using 0.1gram/litre (g/L) of tricaine methanesulfonate; measured for weight (g), length (fork length, mm). A minimum recovery period of one hour was allowed before fish were released at the tagging site.

**Fig. 1.**
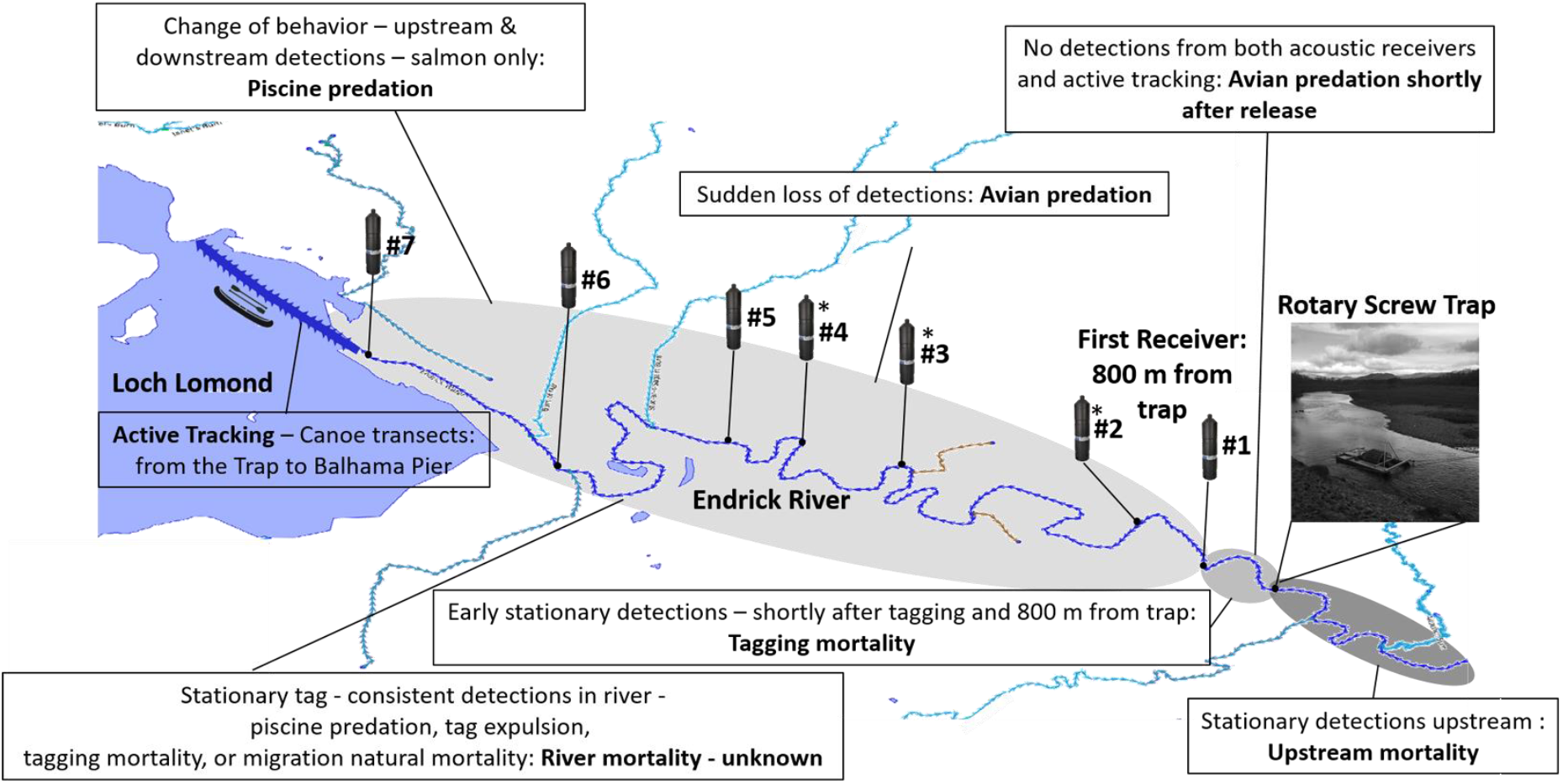
Schematic of the study area and the combined passive and active tracking methodology developed to monitor Atlantic salmon and sea trout smolt migration. The passive acoustic design includes four VR2W receivers initially installed between March 24^th^ and April 14^th^ 2020 (that is prior to fish tagging). Three additional receivers were installed on May 21^th^ 2020, which are represented by *. The active tracking design consist of a canoe transect commencing ∼200 m above the rotary screw trap (and the release site) down river to Balhama Pier, beyond the mouth of the river, using a VR100 portable acoustic reciever. The River Endrick is represented in dark blue. The inflows to the River Endrick are represented in light blue, whereas the outflows are in yellow. Inferences drawn from the patterns of detections (or the lack of detections) from the combined active and passive accoustic receivers led to six plausable migration outcomes for each fish (beside the “unknown”). These inferences are presented in Table 1 and A1.

### Passive Acoustic Tracking Design

To assess smolt migration through the River Endrick, four 69 kHz acoustic receivers (VEMCO VR2W and VR2Tx VEMCO Ltd.) were deployed in the river between March 24^th^ and April 14^th^ 2020. The first receiver was located 800 m downstream from the trapping/release zone and the last receiver was placed at the mouth of river, to detect smolts entering Loch Lomond (#1, 5, 6, and 7; Fig. 1). On May 21^st^ 2020, three additional acoustic receivers (VR2Tx; VEMCO Ltd.) were deployed in the river when the environmental conditions in the river (i.e., the lack of precipitation) resulted into a total obstruction of the smolt migration due to an exposed sandbank at the junction of the river mouth and Loch Lomond. These three additional acoustic receivers (#2-4) were deployed in the mid-section of the river to expand detection coverage from the trapping zone to the mouth of the river (Fig. 1). All receivers were attached to a steel bar welded onto a 20kg weight, which was then anchored to the shore with a rope or chain (Fig. A1).

### Active Acoustic Tracking Design

The active tracking programme consisted of a number of transects of 13.5 km, undertaken by canoe that began ∼200 m upstream of the rotary screw trap, downstream into Loch Lomond (Balhama Pier; 56 ° 08’ 36’’N; 004 ° 54’ 79’’ W). Tags were detected during the passage downstream using a VR100 receiver (VEMCO Ltd.; normal filter setting), designed for manual tracking from small boats, coupled with an omni-directional hydrophone which was attached to the canoe. The operator could adjust the length of the cable to prevent the trailing hydrophone hitting the substrate. Canoe transects were completed daily, beginning three days after tagging commenced, for seven days (that is the April 16^th^ to the 23^rd^ 2020) and every two weeks after that (i.e., April 24^th^ to the June 6^th^ 2020), for an overall total of 10 active tracking surveys. The last canoe transect, completed on June 6^th^ 2020, was started further upstream near a waterfall that acts as a natural barrier for smolts and parr upstream movements. This barrier is ∼ 3.7 km upstream of the release site, resulting in a transect of ∼17. 2 km. This longer transect ensured coverage of all possible upstream sites into which fish may have moved and thus reduced the possibility of undetected tags (see Table 1).

**Table 1.**
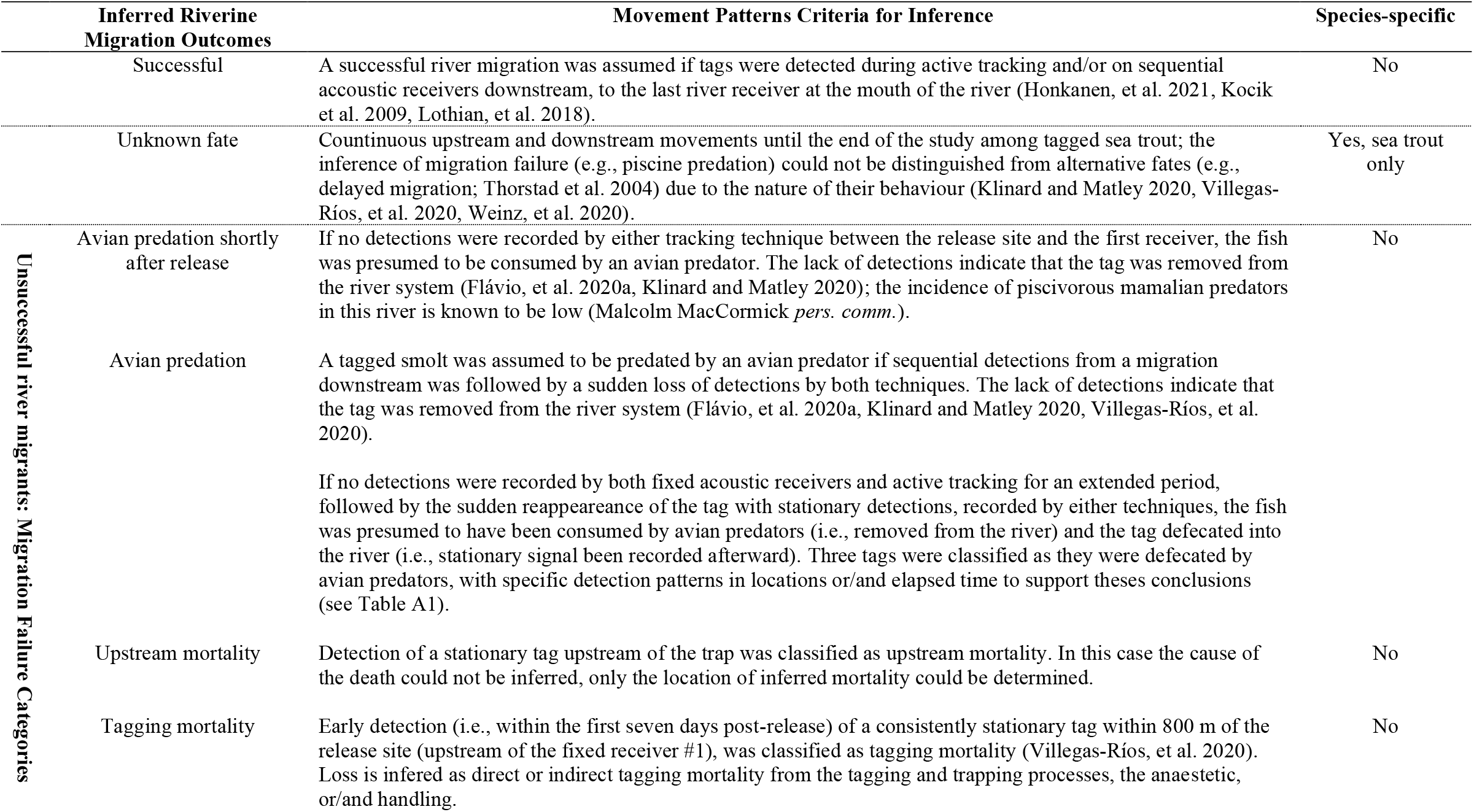
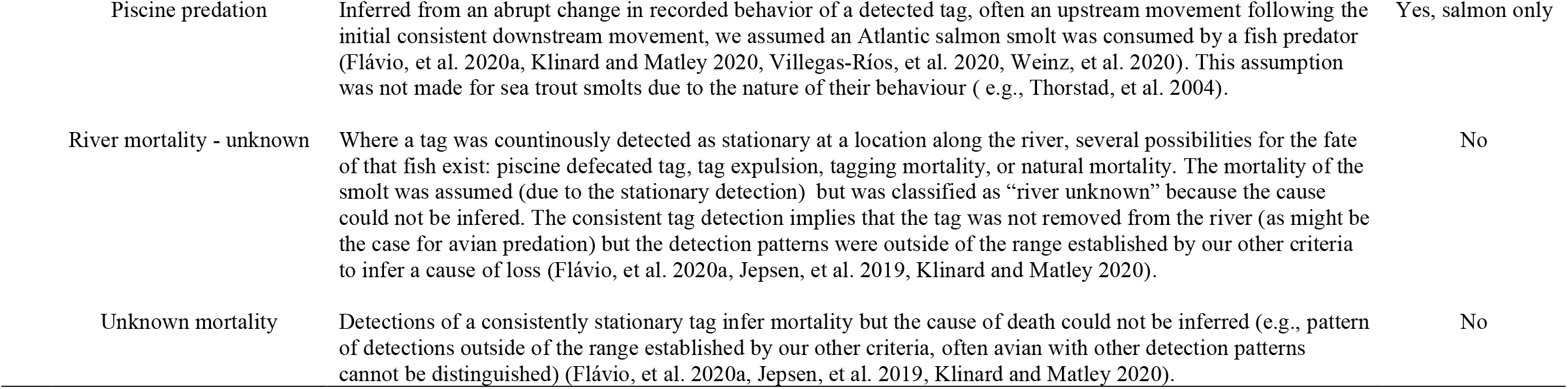
The fate of each tagged fish was inferred following postulations used in other recent studies (for example, see Flávio, et al. 2020a, Gerber, et al. 2017, Klinard and Matley 2020, Villegas-Ríos, et al. 2020, Weinz, et al. 2020), with some modifications for the local study area. Riverine migration outcomes from tag detection patterns were inferred as: successful river migrant, unknown fate, and unsuccessful river migrants that were classified into seven migration failure categories. A detailed movement pattern and justification of our criteria for each fish inferred as dead or as an unknown fate is provided in table A1.

### Detection efficiency

All data were initially compiled in Vemco VUE software and all analyses were conducted in R version 3.5.0 (R Core Team 2016), except if stated otherwise. For calculating detection efficiency of fixed receivers (with the exception of the last receiver downstream), fish detections at a downstream receiver were compared to those from the next upstream receiver, i.e. all fish detected at receiver #2 must have passed receiver #1, providing a detection efficiency measure for receiver #1. Fixed receivers #1, #5 and #6 were included in this analysis (due to their earlier deployment, receiver #2-4 were excluded from this assessment). We also calculated the efficiency of fixed receivers by combining passive and active tracking data; this allowed for the detection of fish between receivers, improving the estimates of fixed receiver detection efficiency. Active tracking detection efficiency was estimated by combining data from active tracking and fixed receivers. Fixed receivers were used to assess which tagged individuals remained within the study area (i.e., those that had not exited the river) and the percentage of these individual detected by active tracking were calculated.

### Sources of smolt riverine migration failure

The pattern of detection of individuals from active and passive acoustic tracking were used to infer the fate of each fish. In the process of evaluating this, we relied on two common assumptions. 1) Tag failure (e.g., through early battery discharge or mechanical failure) was assumed to be minimal. Failure of the tags used in this type of study is generally under two percent (H. H. Honkanen, Unpublished data; Newton et al. 2019). 2) Tag expulsion from the fish was assumed to be inconsequential. This study accumulated data from fish over a few days to a few weeks and where it has been examined, tag expulsion is thought to occur over a longer period (i.e., 40 days; Brunsdon et al. 2019, Lacroix et al. 2004). In addition, to infer an ultimate fate to each tagged fish, we followed postulations used in other recent studies (for example, see Flávio et al. 2020a, Gerber et al. 2017, Klinard and Matley 2020, Villegas-Ríos et al. 2020, Weinz et al. 2020), with some modifications for the local study area, to infer nine categories of riverine migration outcomes from tag detection patterns (see Table 1 and A1 for more details). A Chi-squared test (Zar 2010) was used to determine if the of number of migration failures was spatially randomly distributed throughout the River Endrick by dividing the river into 1 km sections.

### Telemetry analyses

Because the data deviated from normality (Shapiro-Wilk; p-values <0.001), the non-parametric Wilcoxon rank tests, implemented by the kruskalmc function from the pgirmess package (Giraudoux 2012) were used for all telemetry analyses. The four fish with fate categorized as ‘unknown’ were excluded from further evaluation and analyses were done separately on salmon and sea trout smolts. The duration of residency events, defined as the time that a fish spent detected at a single receiver, was compared between fish that were successful and unsuccessful river migrants and among receivers. The rate of movement, measured as the detected distance moved over time elapsed between fixed receivers, of all fish was determined and the difference between successful and unsuccessful migrants and among different river sections (between fixed receivers) were tested with a Kruskal-Wallis chi-squared. To look for diurnal patterns of fish movement and any changes in these patterns, we recorded the time of day that a smolt was first detected at a receiver. Although condition factor did not vary between successful and unsuccessful smolts (Lilly et al. 2021), the effects of fork length and tag burden on smolt migration success were assessed.

## Results

### Detection efficiency

Detection efficiency using the fixed receivers #1, #5, and #6 (Fig. 1) was 98.7%, 100%, and 100%, respectively. When active tracking data were added to efficiency measures, fixed receivers #1, #5, and #6 had a detection efficiency of 95.6%, 100%, and 100%, respectively. For active tracking alone, efficiency increased with time, from 38.5% to 95.2% (Table 2). Overall, 93% (53/57) of the successful migrants, 69% (67/97) of the unsuccessful migrants, and 100% (4/4) of the unknown were detected by active tracking.

**Table 2.** Efficiencies of active tracking, fixed receivers and fixed receivers encompassing active tracking data (see methods) in the River Endrick.

|  | **Active tracking efficiency (%)** |  | **Fixed receiver efficiency (%)** |  | **Fixed receiver efficiency (%) - active tracking** |
| --- | --- | --- | --- | --- | --- |
| April 17<sup>th</sup> | 48 | #1 | 98.7 | #1 | 95.6 |
| April 18<sup>th</sup> | 47.4 | #5 | 100 | #5 | 100 |
| April 19<sup>th</sup> | 45.1 | #6 | 100 | #6 | 100 |
| April 20<sup>th</sup> | 38.5 |  |  |  |  |
| April 21<sup>th</sup> | 42.1 |  |  |  |  |
| April 22<sup>th</sup> | 67.5 |  |  |  |  |
| April 23<sup>rd</sup> | 64.9 |  |  |  |  |
| May 8<sup>th</sup> | 82.6 |  |  |  |  |
| May 21<sup>th</sup> | 86 |  |  |  |  |
| June 4<sup>th</sup> | 95.2 |  |  |  |  |

### Sources of smolt riverine migration failure

In our study, 36.1 % of the Atlantic salmon and sea trout smolts from the River Endrick were assumed to have made a successful river migration, that is, the tag was detected having left the River Endrick. Successful river migration for Atlantic salmon was 36.3%, (n=49; Fig. 2) and for sea trout 34.8% (n=8). 61.4 % of the tagged smolts failed to migrate successfully and were inferred as dead (see Table 1 and A1). For Atlantic salmon, failed river migration was 63.7% (n=86) and for sea trout 47.8% (n=11; see Table 1 and A1). Four sea trout smolts (17.4%), had their migration success categorized as unknown fate (see Table 1 and A1).

**Fig. 2.**
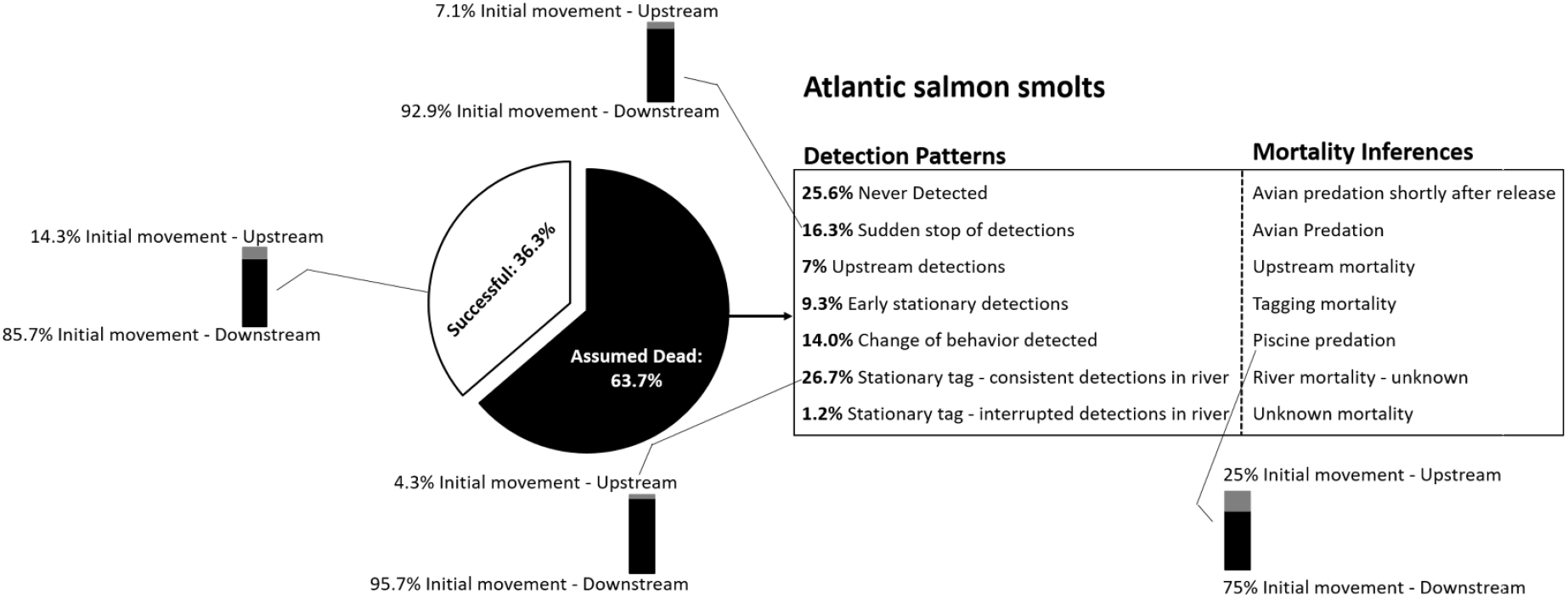
Tags movements of Atlantic salmon recorded with fixed acoustic and active tracking techniques led to the determination of detection patterns that allowed the inference of the fate for each fish (see Table A1 for details). Seven detection patterns were defined to lead to a specific cause of mortality inference (see source of mortality section, table 1 and A1). Initial upstream movement detected were reported with a bar graph.

Of the salmon assumed dead (sea trout excluded due to sample size, see Fig. 2 and Table 1 and A1), 25.6 % were never detected by both methods (n=22; inferred as avian predation shortly after release, near the trap) and 16.3% of smolts suddenly stopped being detected (n=14; inferred as avian predation). Thus, 41.9% of tagged salmon were thought to have been preyed upon by birds. 9.3% of the Atlantic salmon smolts (n=8) were consistently detected at the same place, close to the fish release site and did not move, we infer these to be fish lost as tagging mortality. 14.0 % of the Atlantic salmon smolts (n=12) displayed a pattern of detection indicating a change of behaviour that gave a strong indication of piscine predation. The continuous detection patterns for 26.7% of the smolts (n=23) could not be classified but displayed stationary detections indicative of mortality within the river, which we categorized as river mortality - unknown. 1.2% of Atlantic salmon smolts (n=1, Table A1) were categorized as unknown mortality due to a stationary tag but the detection pattern resulted in a number of possible inferences. 21% Atlantic salmon smolts (n=18) were detected upstream of the release site. Of these, 6 individuals (7%) never initiated a downstream migration, and thus they were classified as upstream mortality. The remaining 12 individuals comprising both successful and unsuccessful migrants, subsequently detected downstream of the release site (see Table A1; Fig. 2).

Finally, mortality inferred events were not randomly distributed along the River Endrick (*X*^2^ =220.54, *df*=13, *P* <0.001; Fig. A2), the highest failure of migration was observed in the first kilometer from the release site.

### Telemetry

Overall, successful river migrants of Atlantic salmon spent a median of 3.99 days (interquartile range (IQR): 2.82-5.22) in the River Endrick, whereas sea trout spent 23.04 days (IQR: 13.98-37.75). For Atlantic salmon, the median duration of all residency events was 1156 seconds (IQR: 544-4109). The duration of residency events was significantly shorter for successful salmon (996 seconds; IQR: 500-2197) compared with unsuccessful river migrant smolts (1926 seconds; IQR: 635-6221; W = 56346, *P* <0.01). A similar but non-significant pattern was found for sea trout, with successful river migrants having shorter residency events (888.5 seconds; IQR: 486.5-2724.3) than unsuccessful river migrants (3431 seconds; IQR: 586-8842; W = 535.5, p = 0.32). The duration of residency events also varied among receivers (Kruskal-Wallis chi-squared = 29.12, *P* <0.01; Table 3), with smolts of Atlantic salmon displaying the longest residency around receiver #1. There were no differences in residency event durations among receivers for sea trout (Kruskal-Wallis chi-squared = 6.34, *P* = 0.096).

**Table 3.** Duration of residency events of smolt specific to each fixed receiver and species in the Endrick.

| **Receiver** | **Species** | **Median duration (sec)** | **IQR (sec)** |
| --- | --- | --- | --- |
| 1 | Salmon | 1916.5 | 610.00 - 7348.00 |
| 1 | Trout | 2791.5 | 577.25 - 9591.25 |
| 2* | Trout | 179.5 | 128.25 - 230.75 |
| 3* | Trout | 407.5 | 274.25 - 540.75 |
| 4* | Trout | 744.5 | 502.75 - 1104.50 |
| 5 | Salmon | 804.5 | 441.00 - 2054.75 |
| 5 | Trout | 659.0 | 418.00 - 940.50 |
| 6 | Salmon | 971.0 | 692.00 - 1224.50 |
| 6 | Trout | 888.5 | 772.00 - 984.25 |
| 7 | Salmon | 1047.0 | 506.00 - 2758.50 |
| 7 | Trout | 1942.5 | 804.75 - 5900.50 |

\*These receivers were deployed late and therefore only include trout movements.

For salmon, the successful river migrants had a higher rate of movement (median: 0.11 m.s^-1^; IQR: 0.05-0.64) than unsuccessful river migrants (0.04 m.s^-1^; IQR: 0.03-0.10; W = 730, *P* <0.01). For sea trout, there was no difference in the rate of movement between successful (median: 0.25 m.s^-1^; IQR: 0.08-0.65) and unsuccessful river migrants (0.09 m/s; IQR: 0.04-0.36 m.s^-1^; W = 61, *P* = 0.53). For both species, rate of movement differed among river sections (salmon: Kruskal-Wallis chi-squared = 105.95, df = 2, *P* <0.01; sea trout: Kruskal-Wallis chi-squared = 16.65, df = 2, *P* <0.01), with the movement rate increasing with distance downstream for both species (i.e., the highest movement rate being at the final section of the river, between receiver #6 and #7). For most of the migration, trout had higher rate of movement than salmon, except for the final section (between receivers #6 and #7) whereas salmon had higher median rate of movement than trout (0.84 and 0.71 m.s^-1^, respectively). During their migration, smolts of Atlantic salmon and sea trout passed the first receiver at all hours of the day, whereas by receivers #6 and #7, most movement was during daytime hours (Fig. 3).

**Fig 3.**
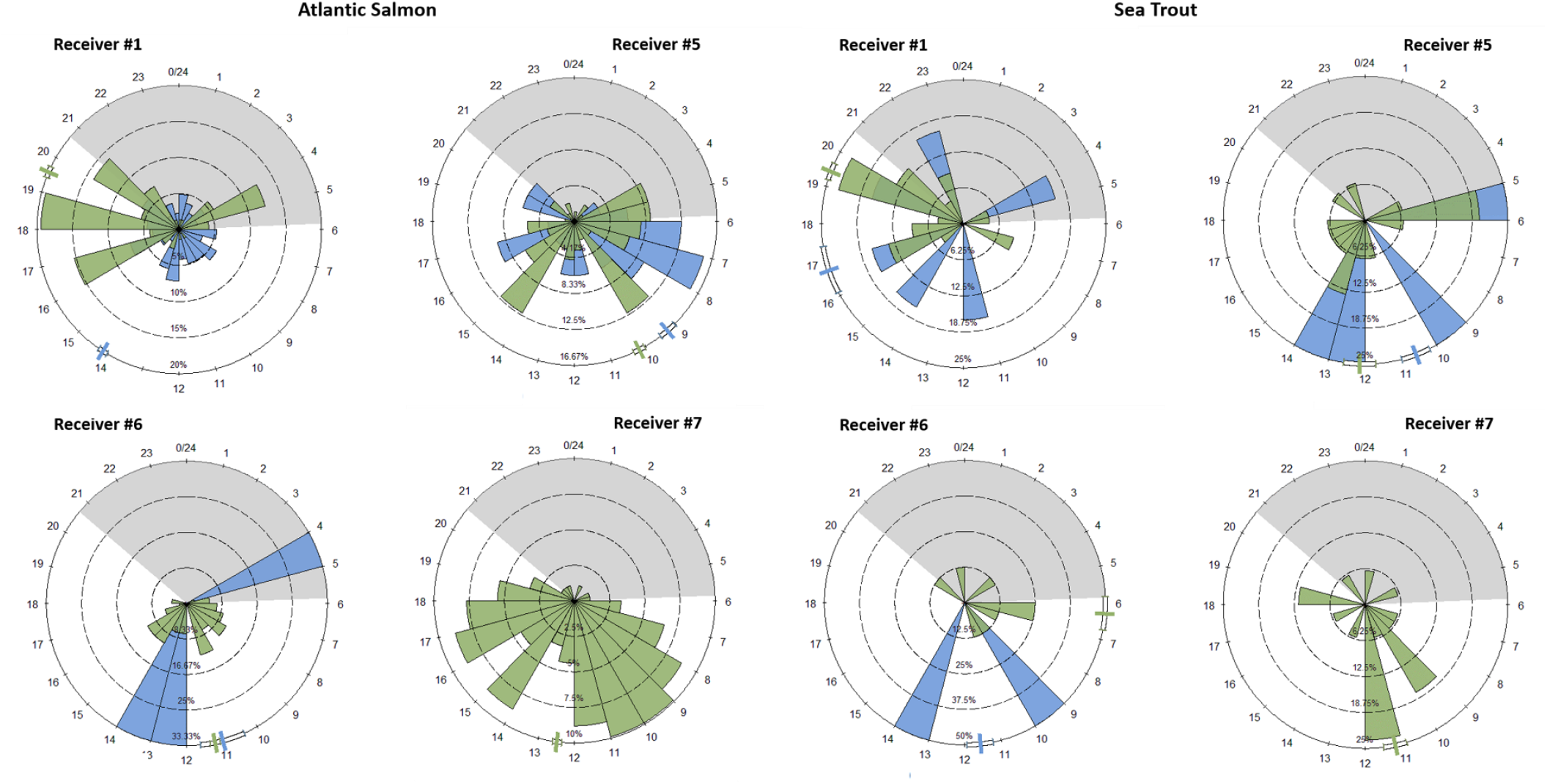
Arrival time at each fixed receiver in the River Endrick showing when Atlantic salmon and sea trout smolts are moving and whether the pattern changes during their migration downstream. Successful (green) and unsuccessful (blue) river migrant smolts are represented at each receiver and each array’s bar sum to 100%. The coloured markers in the outer circle indicate the mean value respectively to the coloured category (i.e., successful and unsuccessful river migrants). The shaded portion of the circle shows the average sunset-to-sunrise hours for April.

Although successful river migrants had a slightly larger fork-length for both species (salmon: 145.3 ± 14.5 mm; trout: 169.6 ± 31.1 mm) than unsuccessful ones (salmon: 141.6 ± 8.9 mm; trout: 168.9 ± 36.3 mm), these size differences were not significant for either species (salmon: W = 1782, *P* = 0.14; trout: W = 41, *P* = 0.87). Additionally, there were no significant differences in tag burden between successful river migrants (salmon: 5.60 ± 1.2%; sea trout: 4.04 ± 1.9%) and unsuccessful river migrants (salmon 5.72 ± 1.0%; trout: 4.05 ± 1.6%; salmon: W = 2156, *P* = 0.82; sea trout: W = 44, *P* = 1).

## Discussion

We provided evidence that the success of river migration by smolts in the River Endrick was low with only 36% of tagged smolts exiting the mouth of the river. This seems to be a trend emerging across rivers in Scotland (Adams, Unpublished data; Honkanen et al. 2021, Lothian et al. 2018) but also elsewhere (Flávio, et al. 2020a, Flávio et al. 2020b, Gibson et al. 2015, Jepsen, et al. 2019) and other salmonid species (Welch et al. 2021, Wilson et al. 2021). Undeniably, the process of smolting is a critical phase in the life-cycle of anadromous salmonids and has been associated with substantial mortality rates (Clark et al. 2016, Halfyard, et al. 2012, Thorstad et al. 2012). Smolt survival can start to be classified as poor when less than 70% of the smolts reach the mouth of the river (Furey et al. 2016), but when riverine survival is lower than half the population, it can have drastic long-standing effects with few adults returning to spawn (Thorstad et al. 2012). Accordingly, one of the pressing requirements to better understand anadromous fish population dynamics is to be able to identify the location, the cause, and the rate of mortality that smolts are facing during their first seaward migration. Here, we produced a framework combining spatial and temporal measures of smolt riverine migration to detect and estimate mortality events using acoustic telemetry. By combining active and passive tracking techniques, we were able to refine the spatiotemporal pattern detection scale, which allowed inferences to be made about mortality location, causes, and rates that would not have been possible using fixed position acoustic receivers solely. The lack of ability to detect fish between two fixed receivers introduces uncertainties that can prevent reasonable inferences from being drawn.

Despite the importance of the smolting phase in anadromous salmonids, the migration from freshwater to marine habitats has been generally less intensively studied than adult spawning migration (Drenner, et al. 2012, Furey et al. 2020). The spatial and temporal patterns of migration failure (e.g., mortality rates, and location) during the smolt riverine migration have the potential to yield important insights into population bottlenecks for anadromous salmonids. In this study we demonstrated that during their initial riverine transitional phase, smolts were particularly vulnerable to predators whilst migrating through that high-risk landscape (Ward and Hvidsten 2011, Ward, et al. 2008). The main cause of mortality of tagged smolts in this study were interpreted as avian predation; mammalian predators is thought to be low in this study area), however, piscine predation was also substantive. The most commonly observed avian piscine species on the River Endrick during this study were goosanders (*Mergus merganser*), grey heron (*Ardea cinerea*), and osprey (*Pandion haliaetus*), whereas the most common aquatic piscine predators in this area will likely be pike (*Esox lucius*) and brown trout (Adams, Unpublished data). Predation can be a major driver of population dynamics and demography (Jonsson et al. 2017, Payton et al. 2020) by exerting a strong selective pressure (Ward and Hvidsten 2011), but other factors can act concomitantly (e.g., environmental factors, hydropower, etc.; Lothian, et al. 2018, Thorstad, et al. 2012). Our finding that avian predation rates were particularly high in the immediate vicinity of the release site, suggests that there was a concomitant interaction between the effects of trapping/tagging and vulnerability to predation for smolts. Thus, the high percentage of individuals that were never detected by either method after release could be a result of either: 1) a learned response by avian predators to the numerical increase in prey near the release site (grey herons (*Ardea cinerea*) and goosanders (*Mergus merganser*) were regularly observed close to the release site; pers. comm.; Flávio, et al. 2020a, Ward, et al. 2008, Wargo Rub and Sandford 2020) and/or 2) individuals were more susceptible to predation following tagging (e.g., trapping, anaesthesia, fish handling; higher residency were observed near the trap; Thedinga et al. 1994). Since the fish were released during the day, it is possible that we induced an experimental artifact by releasing vulnerable fish at a time when visually oriented predators, like birds, are particularly effective. In a previous study in a different Scottish river there was no difference of survival of smolts release during the day vs. night (C.E Adams; unpublished data). Nevertheless, other explanations independent of trapping/tagging induced effects, such as a naturally high predation level in that specific area, cannot be ruled out.

A contentious point of using telemetry has always been to estimate the direct effect of tagging on smolt fate, which has been mostly tested experimentally or using models, but rarely in the wild (Hueter et al. 2006, Klinard and Matley 2020, Newton et al. 2016, Vollset et al. 2020). This study suggests some tagging-induced mortality, but it was unrelated to tag burden. The varying skills of the tagger when inserting tags surgically have been shown to have an effect on the outcome of the tagging process (Cooke et al. 2003) but environmental factors, such as elevated water temperature, can also have an effect on tagging outcomes (Brownscombe, et al. 2019). One way or the other, this result highlights the urgent need to determine direct and indirect effects of surgically implanted tags in telemetry studies to ensure adequate interpretation of the results.

Tagged smolts showed migratory behavioral patterns that were both similar and distinct to other migratory salmonid populations experiencing high predation risks. Residence period and rate of movement of smolts while migrating indicated that a shorter time spent within the river was related to migration success, corroborating the notion that the River Endrick is a high-risk landscape for salmonid juveniles (Furey, et al. 2016, Furey et al. 2021, Honkanen et al. 2018). In this study we did not directly examine the physiological status of migrating smolts. Condition factor might arguably provide an indication of gross physiological condition. However, in this study there was no difference in condition factor between successful and unsuccessful river migrants (Lilly, et al. 2021), which suggest that the process of smolting was well developed in all individuals used in the study and that overall body condition was not a determinant of migration success. We cannot however rule out other physiological drivers of river migration success from this study (McCormick et al. 1999, Thorstad, et al. 2012).

Despite the evidence of high predation pressure found in the river, smolts travelled during the day, especially as they moved further downstream. Typically, smolts travel during the night as a behavioural predation risk-reduction mechanism (Flávio, et al. 2020b, Furey, et al. 2016, Ibbotson et al. 2006). Diurnal travelling is counterintuitive to predation avoidance, particularly in a river with visual predators, and thus migration during the night should lead to lower mortality. However other studies have also found a diurnal pattern (Fängstam et al. 1993, Thorstad, et al. 2012), indicating the occurrence of daytime migration could be related to other factors (e.g., the predator community of a river, latitude, and how early or late in the period of the smolt migration period the fish is).

Upstream movements are considered unlikely for migrating Atlantic salmon smolts and downstream movements are assumed to be ubiquitous (Gauld et al. 2013, Lothian, et al. 2018). Yet, movements of the smolts in the River Endrick offered a diverse range of upstream movements, with a variation in time and final direction (downstream or upstream). Salmon telemetry studies rarely link upstream movements to tag effects or handling and to mortality events occurring upstream after tagging (Frank et al. 2009). These complex behaviours could also be induced from exploratory movements (Keefer et al. 2008), seeking alternative routes, waiting for appropriate conditions (Holbrook et al. 2009), disorientation in certain hydraulic conditions (Honkanen, et al. 2021), or varying sensitivity in distinct migratory phases (Mäkinen et al. 2000). The amount of upstream movement and mortality found in this study highlight the importance to monitoring these movements to ensure adequate interpretation of the results of telemetry studies.

### Important considerations

A few caveats should be noted that could alter interpretations in this study. Despite our best efforts, some uncertainties persisted when mortality sources and rates were untangled and inferred. For example, a number of uncertainties are related to the detection patterns of “never been detected” or “sudden stop in detections”, which were inferred as avian predation. Other factors could induce these patterns, i.e., undetected long-distance upstream movements (no fixed receivers were positioned upstream), mammal predation, and/or tag malfunctioning/expulsion that could result in an overestimation of the avian predation. Although bias in the precision in estimating the proportion of fish in any inferred mortality category is probable; regardless of the uncertainties in our estimations, the main point remaining here is that by far the most substantive detection pattern points to a high percentage of avian predation events.

Our result bias can also be related to underestimating mortality events, which is likely to occur in the delineation of piscine predation events. Due to the difficulty of characterizing a change of behavior, we included only the most obvious patterns from our detections. Most likely, a number of piscine predation mortality events were classified as river mortality – unknown because of the difficulty of distinguishing predator from prey behaviors (vice versa is also possible, a prey behavior could have been quantified as a predator behavior, although less likely due to the strong unidirectional downstream migration patterns displayed by Atlantic salmon). There are also uncertainties associated with the location and timing of these piscine predation events (i.e., where and when the smolt was predated). The use of predation tags (Weinz, et al. 2020), which can detect predation events, coupled with active tracking, could be a valuable method to further refine mortality events of smolts. Finally, due to our strict temporal and spatial assumptions around mortality near the release site, the inferred tagging mortality rate might be an underestimation. In reality, mortality events associated with tagging could take place downstream and upstream of the release site and/or after seven days; in this paper, these events have been classified as river mortality or upstream mortality.

There was a learning curve associated with the active tracking methodology, mostly linked to the disruptive code collisions (i.e., detection interference as a result of several transmissions echoes being heard simultaneously; Binder et al. 2016). Because 53 of 57 successful migrants were detected by active tracking, the increase of detection efficiency over time appears to be linked to the gained experience of handling tag transmission collisions than differences of detection between alive and dead individuals. Even with the poor performance of active tracking at the start of the study, detection efficiency of active tracking improved and refined the scale of detection when those data were combined with fixed receivers.

Despite these caveats, the aim of this paper was to disentangle some aspects of the location, cause, and rate of mortalities occurring and influencing riverine migration success based on several published assumptions with their associated uncertainties (enumerated above). These assumptions are frequently used by studies to target specific mortality causes independently (e.g., avian and piscine predation). Combining spatial and temporal metrics allowed us to attempt to improve the resolution of the tagged smolt fates, while we are acknowledging some uncertainties surrounding identifying location, cause, and rate of mortality events remain. As more studies refine active tracking combined with fixed receivers as a tracking methodology in the riverine environment, assigning location, cause, and rate of morality more precisely is anticipated.

### Conclusion

Active tracking combined with fixed acoustic receivers provided considerable insights into the patterns of migration failure and their causes that could not be identified by relying solely on fixed acoustic receivers. Refining the temporal and spatial scale of the mortality location enabled the causes and rates to be inferred, which we embedded this into a behaviour and movement study. Thus, we provided a case study of a methodology, applicable to any small-moderate riverine system, which should stimulate further use in an emerging research theme and provide insights to help to guide management actions (Flávio, et al. 2020a, Klinard and Matley 2020, Villegas-Ríos, et al. 2020). The extended spatial and temporal characteristics of salmonid migration means that factors acting over long periods and broad geographic scales may all contribute, both cumulatively and synergistically, to the currently depressed populations (Knudsen and McDonald 2020, Thorstad, et al. 2012). Thus, the freshwater and marine survival of anadromous salmonids are known to be intertwined at the population level, although our knowledge of the relationship between the two is limited (McCormick et al. 2009, Sobocinski et al. 2021). Anadromous salmonid marine mortality is considered to be density-independent (Flávio et al. 2019, Jonsson et al. 1998), which implies that the number of juvenile individuals, the smolts, entering seawater will broadly determine the number of adults that return from sea to spawn (McCormick, et al. 2009). Thus, success in the early stages of migration to sea success is likely to have far reaching effects on population dynamics. Considering the major declines of anadromous salmonid worldwide (Chaput 2012), anadromous salmonid populations are of high conservation and scientific relevance, and identifying bottlenecks during their freshwater transition to the marine environment should allow specific management actions to mitigate these losses.

## Acknowledgements

This is a SeaMonitor project funded by the European Union INTERREG VA Programme award number IVA5060. We thank the Loch Lomond Angling Improvement Association, Megan-Rose MacDonald, and Adam Wright for their valuable help with fieldwork and logistics. We thank Matthew Faust for his thoughtful comments on the manuscript.

## Declarations

### Authors’ contributions

LC conceived the study. CEA funded the study. LC, HH, JL, HG, and CEA carried out the field work. LC, HH, JL, MN participated in the data analyses. LC wrote the manuscript. All authors read and approved the final manuscript.

### Availability of data and materials

The datasets supporting the conclusions of this article are included within the article. Raw data will be available on Dryad upon manuscript acceptance.

## Bibliography

Bergé J, Capra H, Pella H, Steig T, Ovidio M, Bultel E, Lamouroux N. 2012. Probability of detection and positioning error of a hydro acoustic telemetry system in a fast-flowing river: Intrinsic and environmental determinants. Fisheries Research. 2012/08/01/;125-126:1–13.

Binder TR, Holbrook CM, Hayden TA, Krueger CC. 2016. Spatial and temporal variation in positioning probability of acoustic telemetry arrays: fine-scale variability and complex interactions. Animal Biotelemetry. 2016/01/28;4:4.

Both A, Duckham M, Laube P, Wark T, Yeoman J. 2012. Decentralized Monitoring of Moving Objects in a Transportation Network Augmented with Checkpoints. The Computer Journal.56:1432–1449.

Brownscombe JW, Lédée EJI, Raby GD, Struthers DP, Gutowsky LFG, Nguyen VM, Young N, Stokesbury MJW, Holbrook CM, Brenden TO, et al. 2019. Conducting and interpreting fish telemetry studies: considerations for researchers and resource managers. Reviews in Fish Biology and Fisheries. 2019/06/01;29:369–400.

Bruneel S, Verhelst P, Reubens J, Baetens JM, Coeck J, Moens T, Goethals P. 2020. Quantifying and reducing epistemic uncertainty of passive acoustic telemetry data from longitudinal aquatic systems. Ecological Informatics. 2020/09/01/;59:101133.

Brunsdon EB, Daniels J, Hanke A, Carr J. 2019. Tag retention and survival of Atlantic salmon (Salmo salar) smolts surgically implanted with dummy acoustic transmitters during the transition from fresh to salt water. ICES J Mar Sci.76:2471–2480.

Carlon R. Tracking tagged fish using a wave glider. Proceedings of the OCEANS 2015-MTS/IEEE Washington; 2015: IEEE.

Chaput G. 2012. Overview of the status of Atlantic salmon (Salmo salar) in the North Atlantic and trends in marine mortality. ICES Journal of Marine Science.69:1538–1548.

Chaput G, Carr J, Daniels J, Tinker S, Jonsen I, Whoriskey F. 2018. Atlantic salmon (Salmo salar) smolt and early post-smolt migration and survival inferred from multi-year and multi-stock acoustic telemetry studies in the Gulf of St. Lawrence, northwest Atlantic. ICES Journal of Marine Science.76:1107–1121.

Clark TD, Furey NB, Rechisky EL, Gale MK, Jeffries KM, Porter AD, Casselman MT, Lotto AG, Patterson DA, Cooke SJ, et al. 2016. Tracking wild sockeye salmon smolts to the ocean reveals distinct regions of nocturnal movement and high mortality. Ecological Applications. 2016/06/01;26:959–978.

Cooke SJ, Graeb BDS, Suski CD, Ostrand KG. 2003. Effects of suture material on incision healing, growth and survival of juvenile largemouth bass implanted with miniature radio transmitters: case study of a novice and experienced fish surgeon. Journal of Fish Biology. 2003/06/01;62:1366–1380.

Cooke SJ, Midwood JD, Thiem JD, Klimley P, Lucas MC, Thorstad EB, Eiler J, Holbrook C, Ebner BC. 2013. Tracking animals in freshwater with electronic tags: past, present and future. Animal Biotelemetry. 2013/05/01;1:5.

Dainys J, Stakėnas S, Gorfine H, Ložys L. 2018. Mortality of silver eels migrating through different types of hydropower turbines in Lithuania. River Research and Applications. 2018/01/01;34:52–59.

Drenner SM, Clark TD, Whitney CK, Martins EG, Cooke SJ, Hinch SG. 2012. A Synthesis of Tagging Studies Examining the Behaviour and Survival of Anadromous Salmonids in Marine Environments. PLOS ONE.7:e31311.

Ennasr O, Holbrook C, Hondorp DW, Krueger CC, Coleman D, Solanki P, Thon J, Tan X. 2020. Characterization of acoustic detection efficiency using a gliding robotic fish as a mobile receiver platform. Animal Biotelemetry. 2020/10/24;8:32.

Fängstam H, Berglund I, Sjöberg M, Lundqvist H. 1993. Effects of size and early sexual maturity on downstream migration during smolting in Baltic samon (Salmo salar). Journal of Fish Biology. 1993/10/01;43:517–529.

Fetterplace LC, Davis AR, Neilson JM, Taylor MD, Knott NA. 2016. Active acoustic tracking suggests that soft sediment fishes can show site attachment: a preliminary assessment of the movement patterns of the blue-spotted flathead (Platycephalus caeruleopunctatus). Animal Biotelemetry. 2016/07/28;4:15.

Flávio H, Aarestrup K, Jepsen N, Koed A. 2019. Naturalised Atlantic salmon smolts are more likely to reach the sea than wild smolts in a lowland fjord. River Research and Applications. 2019/03/01;35:216–223.

Flávio H, Caballero P, Jepsen N, Aarestrup K. 2020a. Atlantic salmon living on the edge: Smolt behaviour and survival during seaward migration in River Minho. Ecology of Freshwater Fish. 2020/08/02;n/a.

Flávio H, Kennedy R, Ensing D, Jepsen N, Aarestrup K. 2020b. Marine mortality in the river? Atlantic salmon smolts under high predation pressure in the last kilometres of a river monitored for stock assessment. Fisheries Management and Ecology. 2020/02/01;27:92–101.

Forseth T, Barlaup BT, Finstad B, Fiske P, Gjøsæter H, Falkegård M, Hindar A, Mo TA, Rikardsen AH, Thorstad EB, et al. 2017. The major threats to Atlantic salmon in Norway. ICES Journal of Marine Science.74:1496–1513.

Frank HJ, Mather ME, Smith JM, Muth RM, Finn JT, McCormick SD. 2009. What is “fallback”?: metrics needed to assess telemetry tag effects on anadromous fish behavior. Hydrobiologia. 2009/11/01;635:237–249.

Furey NB, Hinch SG, Bass AL, Middleton CT, Minke-Martin V, Lotto AG. 2016. Predator swamping reduces predation risk during nocturnal migration of juvenile salmon in a high-mortality landscape. Journal of Animal Ecology. 2016/07/01;85:948–959.

Furey NB, Martins EG, Hinch SG. 2020. Migratory salmon smolts exhibit consistent interannual depensatory predator swamping: Effects on telemetry-based survival estimates. Ecology of Freshwater Fish. 2020/06/17;n/a.

Furey NB, Martins EG, Hinch SG. 2021. Migratory salmon smolts exhibit consistent interannual depensatory predator swamping: Effects on telemetry-based survival estimates. Ecology of Freshwater Fish. 2021/01/01;30:18–30.

Gauld NR, Campbell RNB, Lucas MC. 2013. Reduced flow impacts salmonid smolt emigration in a river with low-head weirs. Science of The Total Environment. 2013/08/01/;458-460:435–443.

Gauld NR, Campbell RNB, Lucas MC. 2016. Salmon and sea trout spawning migration in the River Tweed: telemetry-derived insights for management. Hydrobiologia. 2016/03/01;767:111–123.

Gerber KM, Mather ME, Smith JM. 2017. A suite of standard post-tagging evaluation metrics can help assess tag retention for field-based fish telemetry research. Reviews in Fish Biology and Fisheries. 2017/09/01;27:651–664.

Gibson AJF, Halfyard EA, Bradford RG, Stokesbury MJW, Redden AM, Jech JM. 2015. Effects of predation on telemetry-based survival estimates: insights from a study on endangered Atlantic salmon smolts. Canadian Journal of Fisheries and Aquatic Sciences.72:728–741.

Giraudoux P. 2012. pgirmess: Data analysis in ecology. R package version.1:617.

Halfyard EA, Gibson AJF, Ruzzante DE, Stokesbury MJW, Whoriskey FG. 2012. Estuarine survival and migratory behaviour of Atlantic salmon Salmo salar smolts. Journal of Fish Biology. 2012/10/01;81:1626–1645.

Heupel MR, Semmens JM, Hobday AJ. 2006. Automated acoustic tracking of aquatic animals: scales, design and deployment of listening station arrays. Marine and Freshwater Research.57:1–13.

Holbrook CM, Zydlewski J, Gorsky D, Shepard SL, Kinnison MT. 2009. Movements of Prespawn Adult Atlantic Salmon Near Hydroelectric Dams in the Lower Penobscot River, Maine. North American Journal of Fisheries Management. 2009/04/01;29:495–505.

Honkanen HM, Orrell DL, Newton M, McKelvey S, Stephen A, Duguid RA, Adams CE. 2021. The downstream migration success of Atlantic salmon (Salmo salar) smolts through natural and impounded standing waters. Ecological Engineering. 2021/03/01/;161:106161.

Honkanen HM, Rodger JR, Stephen A, Adams K, Freeman J, Adams CE. 2018. Counterintuitive migration patterns by Atlantic salmon Salmo salar smolts in a large lake. Journal of Fish Biology. 2018/07/01;93:159–162.

Hueter RE, Manire CA, Tyminski JP, Hoenig JM, Hepworth DA. 2006. Assessing Mortality of Released or Discarded Fish Using a Logistic Model of Relative Survival Derived from Tagging Data. Transactions of the American Fisheries Society. 2006/03/01;135:500–508.

Hussey NE, Kessel ST, Aarestrup K, Cooke SJ, Cowley PD, Fisk AT, Harcourt RG, Holland KN, Iverson SJ, Kocik JF, et al. 2015. Aquatic animal telemetry: A panoramic window into the underwater world. Science.348:1255642.

Ibbotson AT, Beaumont WRC, Pinder A, Welton S, Ladle M. 2006. Diel migration patterns of Atlantic salmon smolts with particular reference to the absence of crepuscular migration. Ecology of Freshwater Fish. 2006/12/01;15:544–551.

ICES. 2017. Report of the working group on North Atlantic salmon (WGNAS).

ICES. 2018. Report of the Baltic Salmon and Trout Assessment Working Group (WGBAST), 20–28 March 2018, Turku, Finland.:369.

Ices CS. 2011. Report of the working group on North Atlantic salmon. ICES Document CM/ACOM.9.

Jepsen N, Flávio H, Koed A. 2019. The impact of cormorant predation on Atlantic salmon and sea trout smolt survival. Fisheries Management and Ecology.26:183–186.

Jonsson B, Jonsson M, Jonsson N. 2017. Influences of migration phenology on survival are size-dependent in juvenile Atlantic salmon (Salmo salar). Canadian Journal of Zoology. 2017/08/01;95:581–587.

Jonsson N, Jonsson B, Hansen LP. 1998. The relative role of density-dependent and density-independent survival in the life cycle of Atlantic salmon Salmo salar. Journal of Animal Ecology. 1998/09/01;67:751–762.

Keefer ML, Caudill CC, Peery CA, Boggs CT. 2008. Non-direct homing behaviours by adult Chinook salmon in a large, multi-stock river system. Journal of Fish Biology. 2008/01/01;72:27–44.

Keefer ML, Stansell RJ, Tackley SC, Nagy WT, Gibbons KM, Peery CA, Caudill CC. 2012. Use of Radiotelemetry and Direct Observations to Evaluate Sea Lion Predation on Adult Pacific Salmonids at Bonneville Dam. Transactions of the American Fisheries Society. 2012/09/01;141:1236–1251.

Klinard NV, Matley JK. 2020. Living until proven dead: addressing mortality in acoustic telemetry research. Reviews in Fish Biology and Fisheries. 2020/09/01;30:485–499.

Knudsen EE, McDonald D. 2020. Sustainable fisheries management: Pacific salmon: CRC Press.

Kocik J, Hawkes JP, Sheehan TF, Music PA, Beland KF. Assessing estuarine and coastal migration and survival of wild Atlantic salmon smolts from the Narraguagus River, Maine using ultrasonic telemetry. Proceedings of the American Fisheries Society Symposium; 2009.

Kraus RT, Holbrook CM, Vandergoot CS, Stewart TR, Faust MD, Watkinson DA, Charles C, Pegg M, Enders EC, Krueger CC. 2018. Evaluation of acoustic telemetry grids for determining aquatic animal movement and survival. Methods in Ecology and Evolution. 2018/06/01;9:1489–1502.

Lacroix GL, Knox D, McCurdy P. 2004. Effects of Implanted Dummy Acoustic Transmitters on Juvenile Atlantic Salmon. Transactions of the American Fisheries Society. 2004/01/01;133:211–220.

Leander J, Klaminder J, Jonsson M, Brodin T, Leonardsson K, Hellström G. 2019. The old and the new: evaluating performance of acoustic telemetry systems in tracking migrating Atlantic salmon (Salmo salar) smolt and European eel (Anguilla anguilla) around hydropower facilities. Canadian Journal of Fisheries and Aquatic Sciences. 2020/01/01;77:177–187.

Lilly J, Honkanen HM, McCallum JM, Newton M, Bailey DM, Adams CE. 2021. Combining acoustic telemetry with a mechanistic model to investigate characteristics unique to successful Atlantic salmon smolt migrants through a standing body of water. Environmental Biology of Fishes. 2021/10/23.

Lothian AJ, Newton M, Barry J, Walters M, Miller RC, Adams CE. 2018. Migration pathways, speed and mortality of Atlantic salmon (Salmo salar) smolts in a Scottish river and the near-shore coastal marine environment. Ecology of Freshwater Fish. 2018/04/01;27:549–558.

Mäkinen TS, Niemelä E, Moen K, Lindström R. 2000. Behaviour of gill-net and rod-captured Atlantic salmon (Salmo salar L.) during upstream migration and following radio tagging. Fisheries Research. 2000/03/01/;45:117–127.

McCormick SD, Lerner DT, Monette MY, Nieves-Puigdoller K, Kelly JT, Björnsson BT. Taking it with you when you go: how perturbations to the freshwater environment, including temperature, dams, and contaminants, affect marine survival of salmon. Proceedings of the American Fisheries Society Symposium; 2009:Citeseer.

McMichael GA, Eppard MB, Carlson TJ, Carter JA, Ebberts BD, Brown RS, Weiland M, Ploskey GR, Harnish RA, Deng ZD. 2010. The Juvenile Salmon Acoustic Telemetry System: A New Tool. Fisheries.35:9–22.

Newton M, Barry J, Dodd JA, Lucas MC, Boylan P, Adams CE. 2016. Does size matter? A test of size-specific mortality in Atlantic salmon Salmo salar smolts tagged with acoustic transmitters. J Fish Biol. 2016/09/01;89:1641–1650.

Newton M, Barry J, Dodd JA, Lucas MC, Boylan P, Adams CE. 2019. A test of the cumulative effect of river weirs on downstream migration success, speed and mortality of Atlantic salmon (Salmo salar) smolts: An empirical study. Ecology of Freshwater Fish. 2019/01/01;28:176–186.

Parrish DL, Behnke RJ, Gephard SR, McCormick SD, Reeves GH. 1998. Why aren’t there more Atlantic salmon (Salmo salar)? Canadian Journal of Fisheries and Aquatic Sciences. 1998/01/01;55:281–287.

Payton Q, Evans AF, Hostetter NJ, Roby DD, Cramer B, Collis K. 2020. Measuring the additive effects of predation on prey survival across spatial scales. Ecological Applications. 2020/12/01;30:e02193.

R: A language and environment for statistical computing Vienna, Austria: R Foundation for Statistical Computing. Available from http://www.R-project.org/

Roy R, Beguin J, Argillier C, Tissot L, Smith F, Smedbol S, De-Oliveira E. 2014. Testing the VEMCO Positioning System: spatial distribution of the probability of location and the positioning error in a reservoir. Animal Biotelemetry. 2014/01/06;2:1.

Schwinn M, Aarestrup K, Baktoft H, Koed A. 2017. Survival of Migrating Sea Trout (Salmo trutta) Smolts During Their Passage of an Artificial Lake in a Danish Lowland Stream. River Research and Applications. 2017/05/01;33:558–566.

Sobocinski KL, Greene CM, Anderson JH, Kendall NW, Schmidt MW, Zimmerman MS, Kemp IM, Kim S, Ruff CP. 2021. A hypothesis-driven statistical approach for identifying ecosystem indicators of coho and Chinook salmon marine survival. Ecological Indicators. 2021/05/01/;124:107403.

Thedinga JF, Murphy ML, Johnson SW, Lorenz JM, Koski KV. 1994. Determination of Salmonid Smolt Yield with Rotary-Screw Traps in the Situk River, Alaska, to Predict Effects of Glacial Flooding. North American Journal of Fisheries Management. 1994/11/01;14:837–851.

Thorstad E, kland F, Finstad B, Sivertsgrd R, Bjorn P, McKinleyd R. 2004. Migration speeds and orientation of Atlantic salmon and sea trout post-smolts in a Norwegian fjord system. Environmental Biology of Fishes. 2004/11/01;71:305–311.

Thorstad EB, Whoriskey F, Uglem I, Moore A, Rikardsen AH, Finstad B. 2012. A critical life stage of the Atlantic salmon Salmo salar: behaviour and survival during the smolt and initial post-smolt migration. Journal of Fish Biology. 2012/07/01;81:500–542.

Villegas-Ríos D, Freitas C, Moland E, Thorbjørnsen SH, Olsen EM. 2020. Inferring individual fate from aquatic acoustic telemetry data. Methods in Ecology and Evolution.11:1186–1198.

Vollset KW, Lennox RJ, Thorstad EB, Auer S, Bär K, Larsen MH, Mahlum S, Näslund J, Stryhn H, Dohoo I. 2020. Systematic review and meta-analysis of PIT tagging effects on mortality and growth of juvenile salmonids. Reviews in Fish Biology and Fisheries. 2020/12/01;30:553–568.

Ward DM, Hvidsten NA. 2011. Predation: compensation and context dependence. Atlantic salmon ecology.199–220.

Ward DM, Nislow KH, Folt CL. 2008. Predators reverse the direction of density dependence for juvenile salmon mortality. Oecologia.156:515–522.

Wargo Rub AM, Sandford BP. 2020. Evidence of a ‘ dinner bell’ effect from acoustic transmitters in adult Chinook salmon. Marine Ecology Progress Series.641:1–11.

Weinz AA, Matley JK, Klinard NV, Fisk AT, Colborne SF. 2020. Identification of predation events in wild fish using novel acoustic transmitters. Animal Biotelemetry. 2020/08/09;8:28.

Welch DW, Porter AD, Rechisky EL. 2021. A synthesis of the coast-wide decline in survival of West Coast Chinook Salmon (Oncorhynchus tshawytscha, Salmonidae). Fish and Fisheries. 2021/01/01;22:194–211.

Wertheimer RH, Evans AF. 2005. Downstream Passage of Steelhead Kelts through Hydroelectric Dams on the Lower Snake and Columbia Rivers. Transactions of the American Fisheries Society. 2005/07/01;134:853–865.

Wilson KL, Bailey CJ, Davies TD, Moore JW. 2021. Marine and freshwater regime changes impact a community of migratory Pacific salmonids in decline. Global Change Biology. 2021/10/20;n/a.

Zar J. 2010. Biostatistical analysis. Pearson Education New Jersey: Pearson Education Inc

