## Appendix for "The benefits of merging passive and active tracking approaches: new insights into riverine migration by salmonid smolts" (from another version of this preprint: version 1, 2021-07-26)

**Appendix A**

Table A1. Description for each individual tracked during this study that was inferred as dead or as fate unknown (with the exception of fish that never been detected and classified as avian predation shortly after release). The justification for each inference made for the cause of death is provided, according to criteria described in the methods (see Table 1). The furtherest point of the migration distance each smolt reached in the river is given, as well as the method that made the final detection. Finally, all upstream movements are included in this table, including the few succesful individuals that displayed that behaviour.

| **Tag Id** | **Species** | **Fate** | **Final detection** | **Distance (m)** | **Inferred Riverine Migration Outcomes** | **Movement pattern description and justification of inferred outcomes** |
| --- | --- | --- | --- | --- | --- | --- |
| 36498 | Salmon | Dead | Active tracking | -939 | River (Upstream) | This tag was detected once, by active tracking on the June 4th, almost kilometre upstream from the release site. Due to the species and the distance upstream, death was inferred but not the cause. |
| 36502 | Salmon | Dead | Active tracking | 399 | River (Unknown) | This tag was detected only once, on the last active tracking transect (June 4th), close to the release site location. The trap was removed May 27th, which could impact positive active tracking detection (6 new tags were detected at the release site location on June 4th). The cause of the sole detection is unknown, whether it is due to the loss of the tag in river substrate or a pre-upstream movement. No fixed receiver detections were made. |
| 36504 | Salmon | Dead | Active tracking | 122 | River (Unknown) | This tag was detected only once, on the second last active tracking transect (May 21th) close to the release site location. Why this is the sole detection is unknown, whether it is due to the loss of the tag in river substrate or a pre-upstream movement. No fixed receiver detections were made. |
| 36506 | Salmon | Dead | Active tracking | -103 | River (Upstream) | This tag was detected once, by active tracking on the June 4th, almost kilometre upstream from the release site. Due to the species and the distance upstream, death was inferred but not the cause. |
| 36507 | Salmon | Dead | Active tracking | -942 | River (Upstream) | This tag was detected once, by active tracking on the June 4th, almost kilometre upstream from the release site. Due to the species and the distance upstream, death was inferred but not the cause. |
| 36508 | Salmon | Dead | Fixed receiver | 760 | River (Unknown) | This tag stopped moving close to the fixed receiver #1 and it was detected by active tracking there on the May 8th and 21th, and June 4th. The cause of death could not be inferred because of the lack of detection at the start of the study (thus an inference of early mortality due to tagging was not made). |
| 36510 | Salmon | Dead | Active tracking | -107 | River (Upstream) | This tag was detected once, by active tracking on the June 4th, almost kilometre upstream from the release site. Due to the species and the distance upstream, death was inferred but not the cause. |
| 36511 | Salmon | Dead | Active tracking | 46 | River (Unknown) | This tag was detected only once, on the second last active tracking transect (May 21th), close to the release site location. The cause of death could not be inferred because of the lack of detection at the start of the study (thus an inference of early mortality due to tagging was not made). |
| 36513 | Salmon | Dead | Active tracking | 20 | River (Unknown) | This tag was detected only once, on the last active tracking transect (June 4th), close to the release site location. The trap was removed May 27th, which could impact active positively tracking detection (e.g., 6 new tags were detected at the release site location on June 4th). Why this was the sole detection is unknown, whether it is due to the loss of the tag in river substrate or a pre-upstream movement. No fixed receiver detections were recorded. |
| 36515 | Trout | Unknown | Fixed receiver | 760 | Unknown | The tag was only detected on the fixed receiver #1 (6 days, last day was on July 2nd). Active tracking detected the tag only once, on May 21th. As this is a sea trout, it is possible that the individual did not migrate and remained in the system (with downstream-upstream movements). The alternative possibility is that the smolt was predated by a piscine predator. Due to the pattern of detections up to July, we do not think avian predation is possible. |
| 36516 | Salmon | Dead | Active tracking | 328 | River (Tagging mortality) | This tag was detected at the same location (i.e., stationary tag) by active tracking transects, for three or more transects. At least one these transects is within the first week after tagging (i.e., daily transects) suggesting early mortality, which we infer as tagging mortality due to the close location of detections to the release site (≤800 m). |
| 36517 | Trout | Dead | Active tracking | 112 | River (Tagging mortality) | This tag was detected at the same location (i.e., stationary tag) by active tracking transects, for three or more transects. At least one these transects is within the first week after tagging (i.e., daily transects) suggesting early mortality, which we infer as tagging mortality due to the close location of detections to the release site (≤800 m). |
| 36518 | Salmon | Dead | Fixed receiver | 760 | Avian predation | This tag was tagged on April 14th and detected by the fixed receiver #1 on April 16th. The tag was never detected by active tracking or other fixed acoustic receivers, we infer the tag was removed from the river by an avian predator. |
| 36522 | Salmon | Dead | Active tracking | 52 | River (Tagging mortality) | This tag was detected at the same location (i.e., stationary tag) by active tracking transects, for three or more transects. At least one these transects is within the first week after tagging (i.e., daily transects) suggesting early mortality, which we infer as tagging mortality due to the close location of detections to the release site (≤800 m). |
| 36523 | Trout | Dead | Active tracking | 41 | River (Tagging mortality) | This tag was detected at the same location (i.e., stationary tag) by active tracking, for three or more transects. At least one these transects is within the first week after tagging (i.e., daily transects) to conclude to an early mortality, which we will infer tagging mortality due to the close location of detections to the release site (≤800 m). |
| 36525 | Salmon | Dead | Active tracking | 1380 | River (Unknown) | This tag went downstream and it was detected by the fixed receiver #1. Active tracking transects detected it downstream of the fixed receiver #1 on April 17th where it became stationary. The stationary tag indicated the fish was dead but the cause of death could not be inferred. Due to the relative close distance to the release site (and time elapsed), it is unlikely that the tag was expelled. The tag travelled further than the 800 m zone determined to qualify as an early stationary detection and classified as tagging mortality. |
| 36527 | Salmon | Dead | Active tracking | 5196 | River (Unknown) | This tag was first detected upstream. The tag subsequently moved downstream, whereas detections were stationary after 19th April. The stationary tag indicated the fish was dead but the cause of death could not be inferred. |
| 36528 | Salmon | Dead | Active tracking | 37 | River (Tagging mortality) | This tag was detected at the same location (i.e., stationary tag) by active tracking transects, for three or more transects. At least one these transects is within the first week after tagging (i.e., daily transects) suggesting early mortality, which we infer as tagging mortality due to the close location of detections to the release site (≤800 m). |
| 36530 | Trout | Dead | Fixed receiver | 760 | River (Unknown) | This tag was tagged on April 14th but it was detected by fixed receiver #1 on April 27th and on and off until July 2nd. Active tracking transects in May detected the tag twice, at the same location, downstream of the fixed receiver #1. Fixed receiver detections showed gaps of 6 hrs between detections but nothing longer (e.g., we classified these detections as stationary tag). The stationary tag indicated the fish was dead but the cause of death could not be inferred. Due to the relative close distance to the release site and short elapsed time, it is unlikely that the tag was expelled. However, the tag was tracked in a time that does not qualify in our early stationary detections criteria for tagging mortality. |
| 36531 | Salmon | Dead | Fixed receiver | 1377 | River (Unknown) | This tag was not detected by the fixed acoustic receiver #1 but it was detected by the active tracking at the same location over several transects (downstream of the receiver #1). The stationary tag indicated the fish was dead but the cause of death could not be inferred. Due to the proximity to the release site and the time elapsed, it is unlikely that the tag was expelled. The tag travelled further than the 800 m zone determined to qualify as an early stationary detections criteria for tagging mortality. |
| 36532 | Salmon | Dead | Active tracking | -914 | River (Upstream) | This tag was detected once, by active tracking on the June 4th, almost kilometre upstream from the release site. Due to the species and the distance upstream, death was inferred but not the cause. |
| 36533 | Salmon | Dead | Active tracking | 37 | River (Unknown) | Active tracking detected the tag twice, at the same location, in May and June. The cause of the mortality could not be inferred due to the late detections. |
| 36534 | Salmon | Dead | Active tracking | 400 | River (Unknown) | Active tracking detected this tag twice, at the same location, in May and June. The cause of the mortality could not be inferred due to the late detections. |
| 36535 | Salmon | Dead | Active tracking | 1379 | River (Unknown) | This tag was not detected by the fixed acoustic receiver #1 but it was detected by active tracking at the same locations on several transects. The stationary tag indicated the fish was dead but the cause of death could not be inferred. Due to the proximity to the release site and the time elapsed, it is unlikely that the tag was expelled. The tag travelled further than the 800 m zone determined to qualify as an early stationary detections criteria for tagging mortality. |
| 36536 | Salmon | Dead | Active tracking | 40 | River (Tagging mortality) | This tag was detected at the same location (i.e., stationary tag) by active tracking transects, for three or more transects. At least one these transects is within the first week after tagging (i.e., daily transects) suggesting early mortality, which we infer as tagging mortality due to the close location of detections to the release site (≤800 m). |
| 36538 | Salmon | Dead | Active tracking | 146 | River (Tagging mortality) | This tag was detected at the same location (i.e., stationary tag) by active tracking transects, for three or more transects. At least one these transects is within the first week after tagging (i.e., daily transects) suggesting early mortality, which we infer as tagging mortality due to the close location of detections to the release site (≤800 m). |
| 36539 | Salmon | Dead | Active tracking | 178 | Avian predation | This tag was detected once by active tracking, on April 15th (the tagging date) but the tag was never detected again. We infer the tag was removed from the river by an avian predator. |
| 36540 | Trout | Dead | Active tracking | 4240 | Unknown | This tag moved quickly downstream on April 15th, and was detected by active tracking at the same location on April 17-19th. It was not detected for 2 consecutive days, and then detected on April 22-23th, 200 m downstream. The tag was not detected afterward. The stationary tag indicated the tag was dead but the cause of death could not be inferred. There are confounding patterns that are difficult to interpret, which lead to an interpretation of an unknown cause of death: the tag was stationary at the same location (not moving or dead), but then no detection for two days suggests that the tag was potentially removed by avian predation? Even more surprisingly is that the tag was detected again suggesting an avian excreted tag (?), then no more detections indicating that active tracking failed to detect or that tag had been removed from the river by avian predation (?). |
| 36541 | Salmon | Dead | Active tracking | 1160 | Piscine predation | The tag was detected 1160 metres downstream on April 18-19th but then it moved upstream and was detected at the fixed receiver #1 on April 20th. The tag was not detected April 21-23rd. The tag was detected further upstream again and was detected at its final location on May and June transects. Because of the sudden change of behaviour, we infer piscine predation. The stationary tag might be an indication of tag excretion or the predator position itself. |
| 36543 | Salmon | Dead | Active tracking | 241 | Avian predation | This tag was detected for the first time by active tracking on April 18th but it was not detected during five daily transects, from April 19th-April 23th. It was detected on May 8th and 21th, at the same location, but not detected on June 4th. The non-detection of the tag over five consecutive transects is unlikely due to the performance of the active tracking methodology, but strongly suggests that the individual was removed from the river by avian predation. Avian excretion of the tag in the river would explain the detection in May. |
| 36544 | Salmon | Dead | Active tracking | 7462 | Piscine predation | This tag was detected in all active tracking transects, except for the first one. The tag was detected near the fixed receiver #1 on April 18-19th, then moved quickly downstream overnight. It was detected further downstream (8002 m) but then changed behaviour and moved back upstream. On May 8th, this tag had re-moved downstream (near its most downstream location, 8002 m), where it was detected in both May active tracking sessions. On June 4th, it had moved slightly upstream. Because of the sudden change of behaviour, we infer piscine predation. |
| 36546 | Salmon | Dead | Active tracking | Loch Lomond sand bank | Avian predation | This tag was never detected by any fixed receivers. It was detected by active tracking April 23th, on the sand bank in Loch Lomond. Birds were commonly observed there. This strongly indicates avian predation with the tag detected because it was excreted on that location. |
| 36547 | Salmon | Dead | Active tracking | 39 | River (Unknown) | This tag was detected only once, on the last active tracking transect (June 4th), close to the release site location. The trap was removed May 27th, which could impact positive active tracking detection (6 new tags were detected at the release site location on June 4th). The cause of the sole detection is unknown, whether it is due to the loss of the tag in river. No fixed receiver detections were recorded. |
| 36550 | Salmon | Dead | Active tracking | 10535 | River (Unknown) | This tag was detected during every active tracking session, steadily moving downstream. The tag was detected at receiver #2 to receiver #7 on April 20th. Then it stopped moving, and was detected downstream of that fixed receiver on all other active tracking sessions. The stationary tag indicated the fish was dead but the cause of death could not be inferred. Due to the relative time elapsed, it is unlikely that the tag was expelled. |
| 36552 | Salmon | Dead | Active tracking | 47 | River (Tagging mortality) | This tag was detected at the same location (i.e., stationary tag) by active tracking transects, for three or more transects. At least one these transects is within the first week after tagging (i.e., daily transects) suggesting early mortality, which we infer as tagging mortality due to the close location of detections to the release site (≤800 m). |
| 36554 | Trout | Dead | Active tracking | 40 | River (Tagging mortality) | This tag was detected at the same location (i.e., stationary tag) by active tracking transects, for three or more transects. At least one these transects is within the first week after tagging (i.e., daily transects) suggesting early mortality, which we infer as tagging mortality due to the close location of detections to the release site (≤800 m). |
| 36555 | Trout | Dead | Fixed receiver | 760 | Avian predation | This tag was detected once by fixed receiver #1, April 16th (smolt was tagged April 15th) but it was never detected afterwards. We infer the tag was removed from the river by an avian predator. |
| 36557 | Salmon | Dead | Active tracking | 37 | River (Unknown) | This tag was detected only once, on the last active tracking transect (June 4th), close to the release site location. The trap was removed May 27th, which could impact positive active tracking detection (6 new tags were detected at the release site location on June 4th). The cause of the sole detection is unknown, whether it is due to the loss of the tag in river substrate or a pre-upstream movement. No fixed receiver detections were made. No fixed receiver detections were recorded. |
| 36563 | Salmon | Dead | Fixed receiver | 7700 | Avian predation | This tag was detected by active tracking, moving downstream on 17-19th April. The last detection was by the fixed acoustic receiver # 5 on April 20th. No detections after that date. We infer the tag was removed from the river by an avian predator. |
| 36564 | Salmon | Dead | Active tracking | -92 | Avian predation | This tag was detected once, by active tracking on April 17th, upstream of the release site. It was never detected again strongly suggesting that is was removed from the river by an avian predator. |
| 36565 | Salmon | Dead | Active tracking | 3780 | River (Unknown) | This tag was detected steadily moving downstream – both from active tracking and fixed receivers. It was detected on its final (and same) location on May 2th and June 4th. The stationary tag indicated the fish was dead but the cause of death could not be inferred. |
| 36569 | Salmon | Dead | Fixed receiver | 760 | Avian predation | This tag was tagged on April 16th and it was detected by active tracking on April 17th, downstream to the smolt release site. Then, the tag was detected at fixed receiver #1 on overnight 20-21th April. The tag disappears afterwards. We infer the tag was removed from the river by an avian predator. |
| 36570 | Salmon | Dead | Fixed receiver | 760 | Avian predation | This tag was originally detected upstream by the active tracking, but then was detected at the fixed receiver #1. Afterwards, the tag was not detected by both methods. We infer the tag was removed from the river by an avian predator. |
| 36572 | Salmon | Dead | Active tracking | 5184 | Piscine predation | The tag was originally detected upstream by active tracking but then started its downstream migration but remained stationary at this point from 23th April. On May 8th, active tracking detected the tag 270 m upstream, where it remained there for the rest of the study. Because of the sudden change of behaviour, we infer piscine predation. The stationary tag might be an indication of the tag excretion by or the predator position itself. |
| 36573 | Salmon | Dead | Fixed receiver | 10610 | Piscine predation | The tag was originally detected upstream by active tracking at the first pool above the release site, on April 17th, a day after tagging. On April 19th, the tag started its downstream migration, and was detected 5 km downstream. The tag was stationary on April 20th, but moved 200 m downstream on April 21th. On April 22nd, the tag travelled another 5 km downstream, and was also detected by the receiver #6. On April 23rd, the tag had moved 2.5 km upstream. The next active transect on May 8th detected it few kilometres upstream, whereas the tag remained there afterward. Because of the sudden change of behaviour, we infer piscine predation. The stationary tag might be an indication of the tag excretion by or the predator position itself. |
| 36574 | Trout | Unknown | Fixed receiver | NA | NA | The tag was recorded on May 1st at fixed receiver #1 (upstream before that?) and then the tag moved all the way to the fixed receiver #7, reaching the mouth of the river on May 11th. The tag was then recorded between fixed receiver # 6 and #7 for 9 consecutive days. On May 20th, it moved upstream and was recorded by fixed receiver #1 on June 19th. |
| 36580 | Salmon | Dead | Active tracking | 200 | River (Unknown) | This tag was detected May 8th and thereafter by active tracking. The cause of death could not be inferred because of the lack of detection at the start of the study to qualify as an early stationary detections criteria for tagging mortality. |
| 36581 | Salmon | Dead | Active tracking | 1530 | Avian predation | The tag was detected near receiver #5 by active tracking on April 19th, 22th and 23rd, but then there were no detections, from active or fixed receivers. We infer the tag was removed from the river by an avian predator. |
| 36582 | Salmon | Dead | Active tracking | 40 | River (Tagging mortality) | This tag was detected at the same location (i.e., stationary tag) by active tracking transects, for three or more transects. At least one these transects is within the first week after tagging (i.e., daily transects) suggesting early mortality, which we infer as tagging mortality due to the close location of detections to the release site (≤800 m). |
| 36584 | Salmon | Dead | Fixed receiver | 7700 | Avian predation | This tag was never detected by active tracking. The tag migrated quickly downstream and was detected at the fixed receiver #1 the same day it was tagged. On April 20th, it was detected at the fixed receiver #5, but was never detected again. We infer the tag was removed from the river by an avian predator. |
| 36585 | Salmon | Dead | Fixed receiver | 760 | Avian predation | This tag was detected at the fixed receiver #1 on the same evening that it was tagged. Not detected since. There were no detections by either method. We infer the tag was removed from the river by an avian predator. |
| 36586 | Salmon | Dead | Active tracking | 400 | River (Unknown) | This tag was detected only once, on the last active tracking transect (June 4th), close to the release site location. The trap was removed May 27th, which could impact positive active tracking detection (6 tags were detected at the release site location on June 4th). The cause of the sole detection is unknown, whether it is due to the loss of the tag in river substrate or a pre-upstream movement. No fixed receiver detections were made. |
| 36590 | Salmon | Dead | Active tracking | 9688 | River (Unknown) | Detections from both methods showed steady movement downstream, until May 8th, where the tag was detected at the same location thereafter. By May 8th, only 20 days have passed since surgery, tag expulsion is a possibility, but outside of the range established by our assumptions. Thus, because of the stationary tag, we inferred the fish was dead but the cause of death could not be determined. |
| 36592 | trout | Unknown | Fixed receiver | 10070 | Unknown | The tag moved downstream and it was detected near the fixed receiver # 6 on April 23rd at 5pm. Four hours later, the tag was detected upstream at previous fixed acoustic receiver, #5. The smolt was detected there for 10 mins, then it disappeared. The tag is back there the next morning, for about an hour. Then, the tag disappeared but it was back the same evening. A similar pattern was detected on April 25th. The smolt was not detected on April 26th but it was back April 27th morning. On April 27th, the tag moved to fixed receiver #2 to fixed receiver #6 before returning to the previous fixed receiver #5. The same downstream-upstream pattern continues on April 28th. April 28th is the last time it is detected by fixed receivers. On May 8th, active tracking transect detect the smolt 3km upstream. The tag was detected at that location afterward by active tracking. Due to the fact that it is a sea trout, it is possible that this individual did not migrate and remained in the system (with downstream-upstream movements). The alternative possibility is piscine predation. |
| 36593 | Trout | Dead | Fixed receiver | 760 | Avian predation | The tag was tagged on April 18th and was detected at the fixed receiver #1 the next morning (19th April), then it disappears. There are no detections by both techniques. We infer the tag was removed from the river by an avian predator. |
| 36595 | Salmon | Dead | Active tracking | 9896 | Piscine predation | The tag was detected near receiver #2, on April 20-23rd. On May 1st, the tag is detected at the fixed receiver #6. On May 8th and afterward, the tag was detected ~700 meters upstream. Because of the sudden change of behaviour, we infer piscine predation. The stationary tag might be an indication of the tag excretion or the predator position itself. |
| 36596 | Salmon | Dead | Active tracking | 9617 | River (Unknown) | Between April 20-21th, the tag made a long distance movement downstream. Stationary location was recorded on April 22th. Small upstream movement (~130 m) was detected on 23th April but came back on the same location of 21-22th April until the end of the study. The stationary tag indicated the fish was dead but the cause of death could not be inferred. |
| 36599 | Salmon | Dead | Fixed receiver | 7700 | River (Unknown) | This tag displayed fast and steady movement downstream, until April 22th. Next day, a small upstream movement was detected by active tracking and detections remained there until the last transect. It was almost continuously detected by the fixed receiver #5 until July. The stationary tag indicated the fish was dead but the cause of death could not be inferred. |
| 36604 | Salmon | Dead | Active tracking | 6883 | Avian predation | A rapid and long distance movement downstream was detected between April 20-22th but the tag was not detected on April 23th. On May 8th, it was detected at Balmaha boat yard (Loch Lomond), where the tag was detected at that location for the remaining study. This pattern indicates avian predation and suggests that the tag was excreted at the boat yard. |
| 36606 | Trout | Unknown | Active tracking | 1474 | Unknown | This tag displayed constant movements between the release site and receiver #2 during the entire study. The tag was detected by both fixed receivers and active tracking. Due to the fact that it is a sea trout, it is possible the individual did not migrate and remained in the river system (with downstream-upstream movements). The alternative possibility is that the smolt was predated by a piscine predator. The tag was constantly moving up to the end of the study, which reinforces the first explanation of the smolt being alive but not migrating. |
| 36607 | Salmon | Dead | Active tracking | 7700 | Avian predation | A very fast movement downstream was detected in less than 24 hrs, but afterwards, the tag was never detected by both techniques. We infer the tag was removed from the river by an avian predator. |
| 36609 | Salmon | Dead | Passive receiver | 5196 | Piscine predation | The tag was originally detected upstream, on the same day of tagging (April 19th). On April 20th, the tag was detected at the receiver #1. On April 21th, the tag had moved 4.3 kilometres downstream. On April 22nd, the tag moved further downstream and was detected ~100 m downstream from receiver #5. However, the tag moved back upstream; by 23 April it had moved 1 km upstream. The tag changed direction again and moved back downstream to receiver #5. The tag was detected by the fixed receiver #5 on April 25th for 8 hrs (this was the final time it was detected by a fixed receiver). The next active tracking session on May 8th detected the tag 3km upstream from the fixed receiver #5 and was also detected there on 21 May and 4 June. Because of the sudden change of behaviour, we infer piscine predation. |
| 36610 | Salmon | Dead | Active tracking | 7700 | Piscine predation | A rapid long distance movement downstream was determined, where the tag was detected by the fixed receiver #5 on April 21th. Two hours later, active tracking detected the tag upstream. On April 22th, active tracking detected the tag one kilometre further upstream. By the next day, active tracking detected the tag another two kilometre upstream. In May and June transects, the tag was detected back downstream, at the same location, near fixed receiver #5. Because of the sudden change of behaviour, we infer piscine predation. The stationary tag might be an indication of the tag excretion by or the predator position itself. |
| 36611 | Salmon | Dead | Active tracking | 7895 | Piscine predation | An initial rapid and long distance movement downstream was detected. On April 21th, active tracking detected the tag near the fixed receiver #5. On April 22th, active tracking detected the tag one kilometre upstream. The next day, active tracking detected the tag two kilometres upstream. In May and June transects, the tag was detected back downstream, at the same location, near receiver #5. Because of the sudden change of behaviour, we infer piscine predation. The stationary tag might be an indication of tag excretion by or the predator position itself. |
| 36614 | Salmon | Dead | Active tracking | 9973 | Piscine predation | A rapid movement downstream was detected, nearly as far as receiver # 6 by active tracking on April 21th. Then, the tag was detected upstream by the fixed receiver #5. The next day, on April 22th, the tag was detected ~three kilometres upstream by active tracking. On April 23rd, it is back at the fixed receiver #5 where it is detected twice a day (morning and evening). On April 23th, the active tracking detected the tag further downstream, near the receiver #6. Between April 27-28th, the tag moved the same downstream-upstream pattern. Active tracking on May 8th and onward detected the tag back upstream, at the same location. Because of the sudden change of behaviour, we infer piscine predation. The stationary tag might be an indication of the tag excretion by or the predator position itself. |
| 36618 | Salmon | Dead | Active tracking | 7700 | Piscine predation | A rapid and long distance movement downstream was recorded on April 19th. The tag was detected close to the fixed receiver #5 on April 22th. The following day, the tag was detected upstream, thereafter it was detected at the same location through all active transects. Because of the sudden change of behaviour, we infer piscine predation. The stationary tag might be an indication of the tag excretion by or the predator position itself. |
| 36621 | Salmon | Dead | Active tracking | -660 | River (Upstream) | This tag was detected once, by active tracking on the June 4th, almost kilometre upstream from the release site. Due to the species and the distance upstream, death was inferred but not the cause. |
| 36623 | Salmon | Dead | Active tracking | 459 | River (Unknown) | This tag was detected only once, on the last active tracking transect (June 4th), close to the release site location. The trap was removed May 27th, which could impact positive active tracking detection (6 new tags were detected at the release site location on June 4th). The cause of the sole detection is unknown, whether it is due to the loss of the tag in river substrate or a pre-upstream movement. No fixed receiver detections were made. |
| 36664 | Salmon | Dead | Active tracking | 10120 | Piscine predation | A very rapid and large movement downstream was detected on April 21th, the tag was almost near the fixed receiver #6 within 24 hrs. On April 22th, the tag was detected several kilometres upstream. On April 23th, the tag was detected downstream, by the fixed receiver #5. On May 8th, active tracking detected the tag several kilometres upstream, where it was detected at the same location in May 21th and June 4th. Because of the sudden change of behaviour, we infer piscine predation. The stationary tag might be an indication of the tag excretion by or the predator position itself. |
| 36666 | Salmon | Dead | Active tracking | 10068 | Piscine predation | This tag was detected by active tracking on April 22th, near the receiver #6. Next day, on April 23rd, the tag was detected upstream. On May 8th, active tracking detected the tag downstream. It stayed at this same location to May 21th and June 4th. Because of the sudden change of behaviour, we infer piscine predation. The stationary tag might be an indication of the tag excretion by or the predator position itself. |
| 36667 | Salmon | Dead | Fixed receiver | 3720 | Unknown | On April 22th, the tag was detected at 3720 m downstream from the release site. The tag changed direction, migrated upstream and was detected at the fixed receiver #1 on April 22th. It was also detected by fixed receiver #1 on the 25th, 26th, and 30th of April. Then, almost constantly detected until July. The tag was not detected by active tracking on April 23rd, May 8th, and June 4th, but it was detected in May 21th. The stationary position of the tag and the non-detections via fixed receivers suggest avian predation and the excretion of the tag at the fixed receiver #1. However, the alternative possibility of piscine predation cannot be ruled out. |
| 36668 | Salmon | Dead | Fixed receiver | 760 | Avian predation | This tag was detected at the fixed receiver #1 on April 23rd and 24th. Active tracking detected the tag at the fixed receiver #1 on April 23rd. No detections were recorded afterwards from both techniques. We infer the tag was removed from the river by an avian predator. |
| 36671 | Salmon | Dead | Active tracking | 418 | River (Unknown) | The tag was detected only once, on the last active tracking transect (June 4th), close to the release site location. The trap was removed May 27th, which could impact active positively tracking detection (e.g., 6 new tags were detected at the release site location on June 4th). The cause of the sole detection is unknown, whether it is due to sedimentation or a pre-upstream movement. No fixed receiver detections were recorded. |
| 36674 | Salmon | Dead | Active tracking | 9390 | River (Unknown) | The tag was detected near the fixed receiver #1 on April 21th and 23rd. By May 8th, active tracking detected the tag downstream of the fixed receiver #5, where it was detected at that location until June. By May 8th, only 20 days have passed since surgery, tag expulsion is a possibility, but outside of the range established by our assumptions. Thus, because the stationary tag, we inferred the fish was dead but the cause of death could not be determined. |
| 36677 | Salmon | Dead | Active tracking | 618 | River (Tagging mortality) | This tag was detected at the same location (i.e., stationary tag) by active tracking transects, for three or more transects. At least one these transects is within the first week after tagging (i.e., daily transects) suggesting early mortality, which we infer as tagging mortality due to the close location of detections to the release site (≤800 m). |
| 36885 | Trout | Dead | Active tracking | 12330 | River (Unknown) | Originally the tag went upstream before starting downstream movement. Tis tag moved fairly quickly to the fixed receiver #7 on April 27th. However tag returned upstream and was detected at the receiver #5 on April 28th and 30th. Active tracking detected the tag back downstream and it was detected on this same location on 8 May, 21 May and 4 June. |
| 36887 | Trout | Dead | Active tracking | 423 | Avian predation | This tag was detected by active tracking, before the fixed receiver #1, on April 22th and 23th, at the same location. No more detection afterwards from both technics. We infer the tag was removed from the river by an avian predator. |
| 36688 | Trout | Dead | Fixed receiver | 760 | Unknown | The tag was never detected by active tracking. The tag was detected at the fixed receiver #1 in three separate day, on April 21th, April 22th, and April 28th. The lack of detection of all active tracking suggest avian predation and the excretion of the tag near the fixed receiver #1. However, piscine predation and failure from active tracking cannot be ruled out. |

Table A2. Tag date and number of Atlantic salmon smolts and Sea trout smolts within the Endrick catchment.

|  | Species | |
| --- | --- | --- |
| Tag Date | Salmon | Trout |
| April 13^th^ | 19 | 3 |
| April 14^th^ | 11 | 1 |
| April 15^th^ | 16 | 3 |
| April 16^th^ | 13 | 5 |
| April 18^th^ | 27 | 4 |
| April 19^th^ | 26 | 2 |
| April 20^th^ | 23 | 5 |

Table A3. Duration of residency events of smolt specific to each fixed receiver and species in the Endrick.

| **Receiver** | **Species** | **Median duration (sec)** | **IQR (sec)** |
| --- | --- | --- | --- |
| 1 | Salmon | 1916.5 | 610.00 - 7348.00 |
| 1 | Trout | 2791.5 | 577.25 - 9591.25 |
| 2* | Trout | 179.5 | 128.25 - 230.75 |
| 3* | Trout | 407.5 | 274.25 - 540.75 |
| 4* | Trout | 744.5 | 502.75 - 1104.50 |
| 5 | Salmon | 804.5 | 441.00 - 2054.75 |
| 5 | Trout | 659.0 | 418.00 - 940.50 |
| 6 | Salmon | 971.0 | 692.00 - 1224.50 |
| 6 | Trout | 888.5 | 772.00 - 984.25 |
| 7 | Salmon | 1047.0 | 506.00 - 2758.50 |
| 7 | Trout | 1942.5 | 804.75 - 5900.50 |

*These receivers were deployed late and therefore only include trout movements.

Table A4. Eigenvector of linear measurements PCAs, each scaled to its standard deviation, (see Fig. A4) of Atlantic salmon and sea trout (in grey shadow) smolts. Principal components, PC1 and PC2, of Atlantic salmon explained 24.16% and 14.75% of the variance, respectively, whereas PC1 and PC2 of sea trout explained 25.04% and 22.82% of the variance.

| **Morphological Traits** | **Eigenvector 1** | **Eigenvector 2** | **Eigenvector 1** | **Eigenvector 2** |
| --- | --- | --- | --- | --- |
| **Upper jaw** | 0.72 | 0.04 | 0.40 | 0.59 |
| **Eye length** | 0.75 | -0.15 | -0.72 | 0.45 |
| **Snout-eye** | 0.80 | -0.34 | 0.39 | 0.66 |
| **Head length** | 0.83 | -0.13 | 0.15 | 0.91 |
| **Dorsal fin** | 0.26 | 0.51 | -0.33 | 0.67 |
| **Caudal fin** | 0.29 | 0.37 | -0.59 | 0.27 |
| **Peduncle depth** | 0.13 | 0.76 | 0.69 | 0.35 |
| **Peduncle length** | -0.15 | -0.12 | -0.65 | 0.03 |
| **Anal fin** | 0.15 | 0.36 | 0.45 | -0.04 |
| **Pelvic fin** | 0.09 | 0.001 | 0.18 | -0.49 |
| **Pectoral fin** | 0.48 | -0.08 | 0.05 | 0.20 |
| **Body depth** | 0.13 | 0.70 | 0.75 | -0.07 |

Table A5. Results from the Multiple Regression Quadratic Assignment Procedure (MRQAP) for all fixed receivers data, testing for a correlation between the presence of a dyad and their binned fork length (mm) or weight (g), whether they were the same species and whether they had the same capture date.

| Predictor | **All Time** | |
| --- | --- | --- |
|  | Coefficient | *p* |
| Fork length | -0.002 | 0.24 |
| Weight | 0.004 | 0.06 |
| Species | 0.003 | 0.22 |
| Tagging day | 0.008 | 0.003 |

*All predictors were not significantly correlated with co-occurrences during each tag date analysed. The variance explained by the models was extremely low (20-04-18: *R*^2^ = 0.01). All predictors were not significantly correlated with co-occurrences during each tag date analysed.

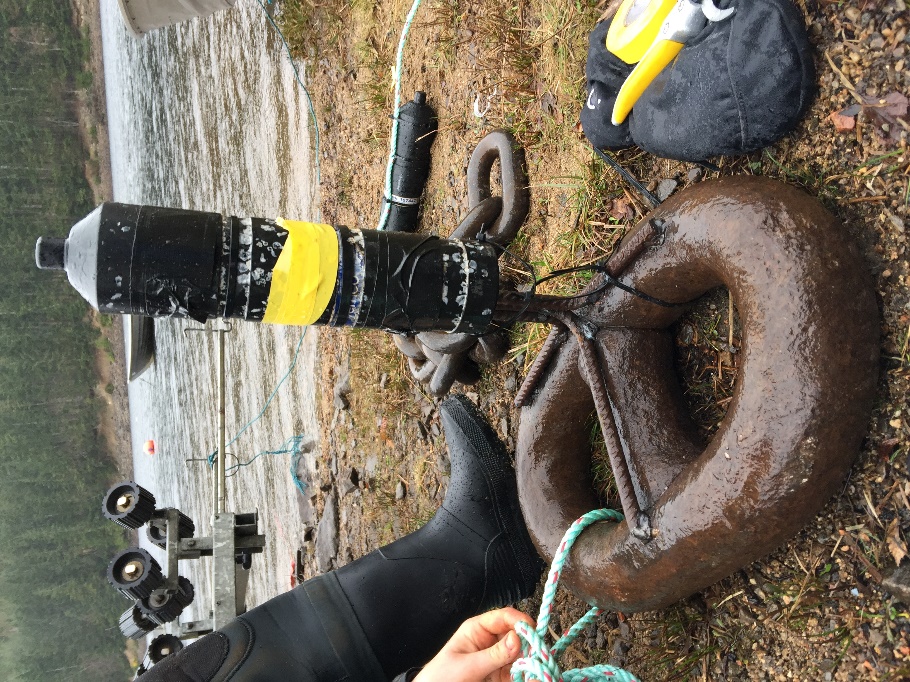

Fig. A1. Depiction of the acoustic receiver set up for the River Endrick, Leven and Clyde Estuary. Receivers were attached to rebar that was welded onto a 20kg weight, weights were then anchored to shore with rope or chain (Photo: Dr. Hannele Honkanen 2019).

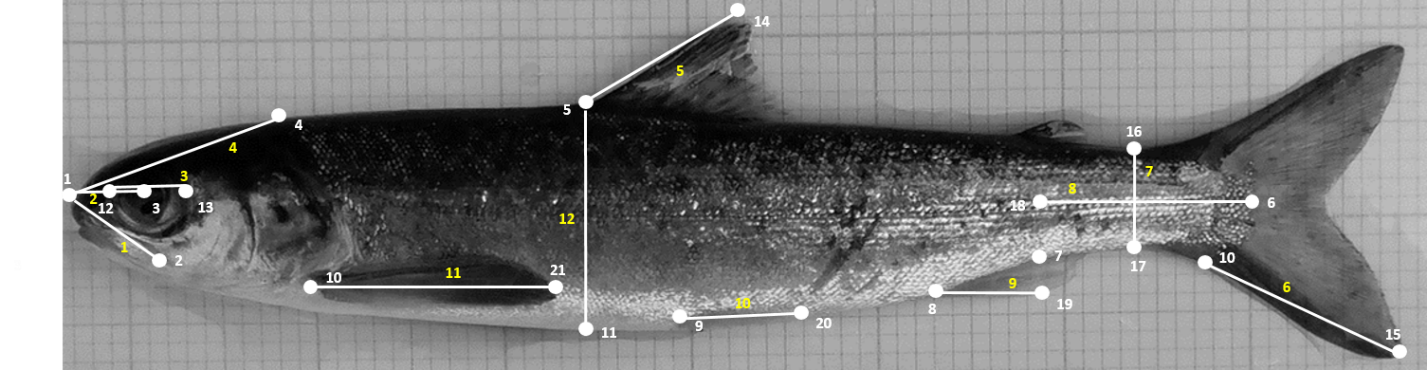

Fig. A2. Landmarks used to measure linear measurements of Atlantic salmon and sea trout from the River Endrick: (1) anterior tip of the snout, (2) posterior tip of maxilla, (3) center of eye, (4) posterior of neurocranium above tip of opercle, (5) dorsal fin anterior insertion, (6) hypural plate midpoint, (7) anal fin anterior insertion, (8) anal fin posterior insertion, (9) pelvic fin insertion, (10) pectoral fin insertion, (11) edge of the ventral surface, positioned directly below the dorsal fin insertion, (12-13) anterior and posterior parts of the eye, (14) dorsal fin tip, (15) caudal fin tip, (16-17) upper and lower points of the least vertical depth of the caudal peduncle, (18) positioned directly above the anal fin anterior insertion, on the lateral line, (19) anal fin tip, (20) pelvic fin tip, and (21) pectoral fin tip. Fin and body length measurements of salmon and sea trout were measured as follow: (1) upper jaw, (2) snout–eye, (3) eye diameter, (4) head length, (5) dorsal fin, (6) caudal fin, (7) caudal peduncle depth, (8) caudal peduncle length (9) anal fin, (10) pelvic fin, (11) pectoral fin, and (12) body depth.

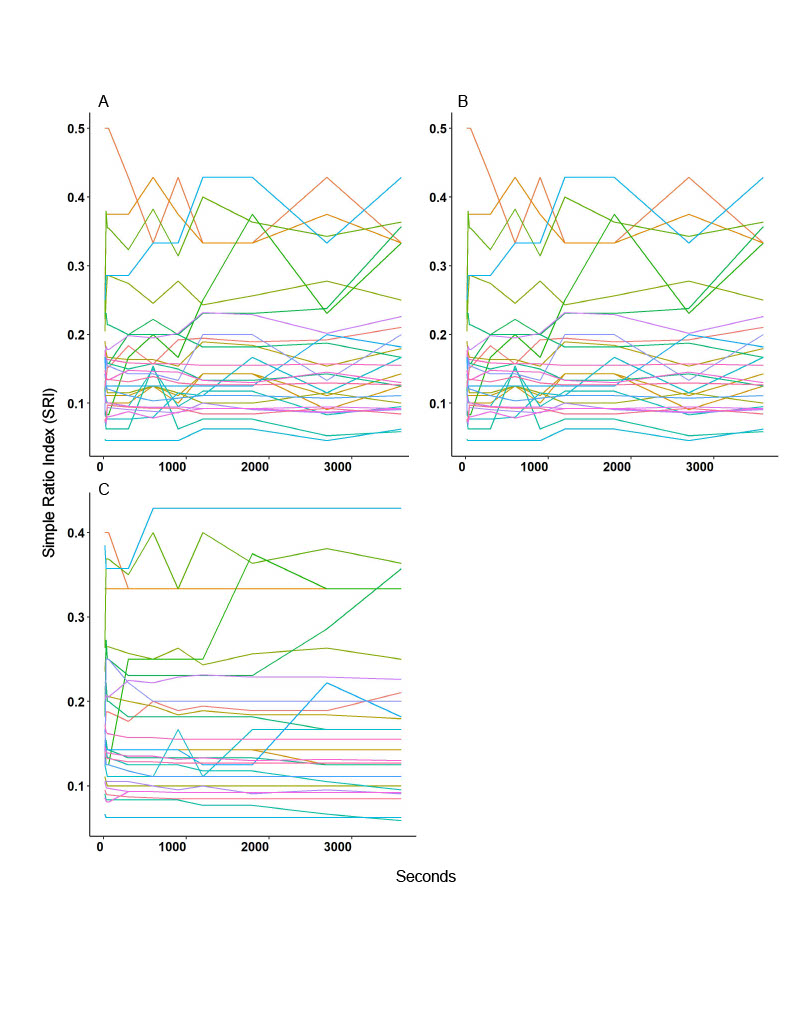

Fig. A3. Sensitivity analyses comparing the differences in strength of the Simple Ratio Index (SRI) during sampling periods of 15-3600 seconds and interval periods of 30 (A), 45 (B) and 60 minutes. Each unique colour represents the SRI for one pair of co-occurring Atlantic salmon and/or sea trout.

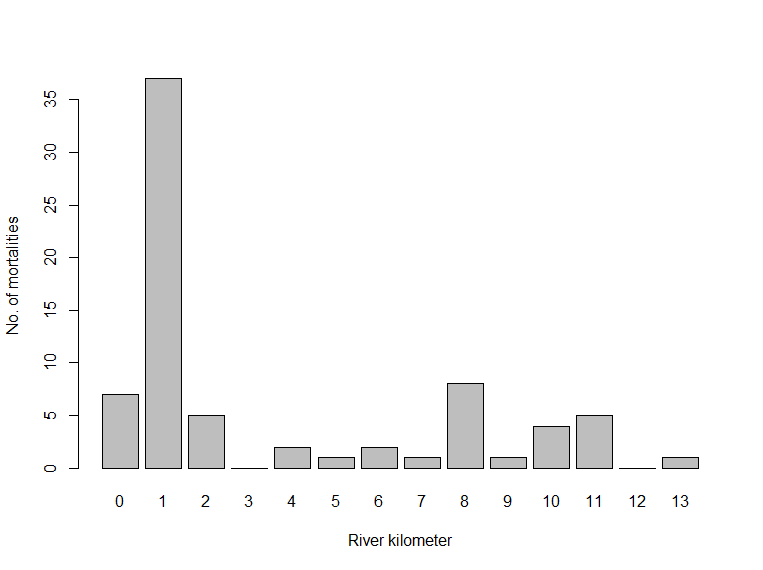

Fig. A4: Spread of mortalities over river distance. 0 is for fish dead upstream from the trap. 1 is for the first kilometer from the smolt trap. River kilometer 13 is the mouth of the river.

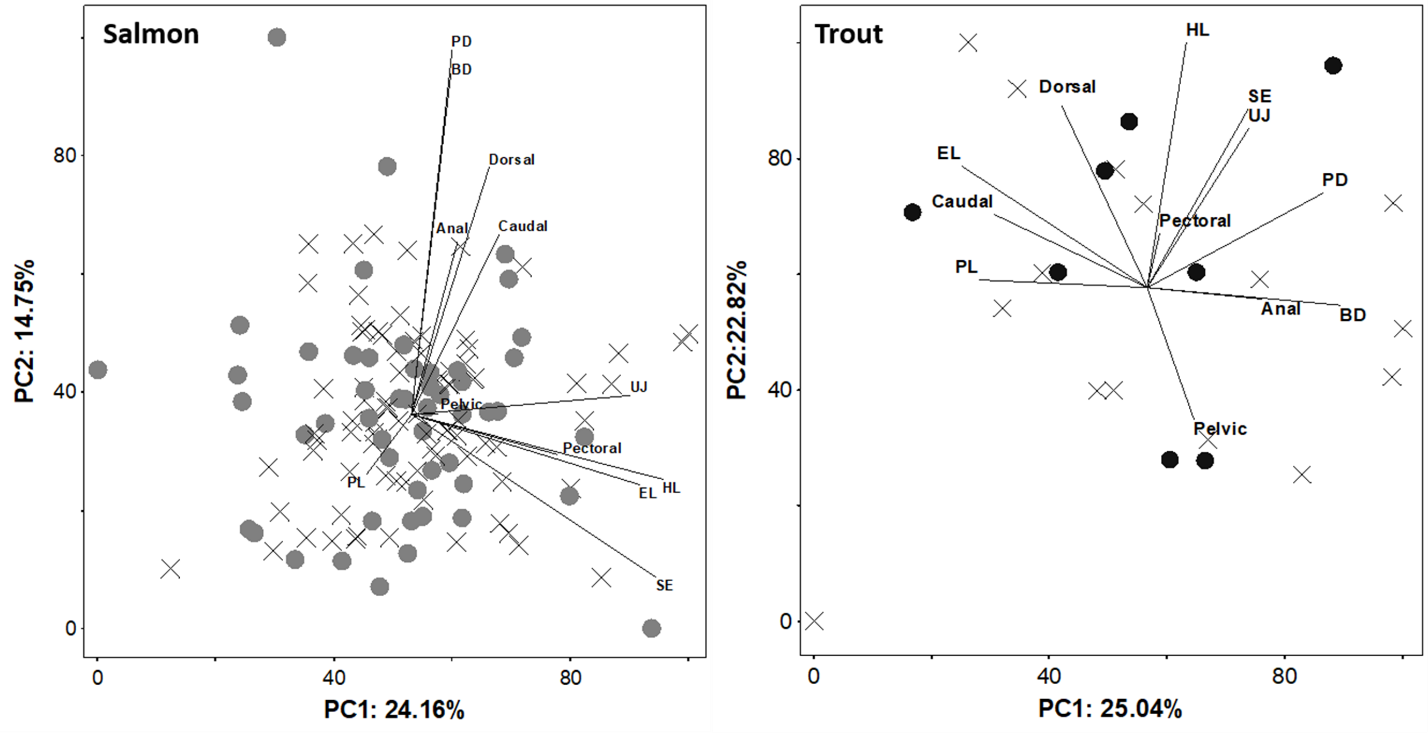

Fig. A5. The PCA ordinations of linear measurements of Atlantic salmon and sea trout. Linear measurement variables are defined in Fig A3 and are represented as follows: UJ = upper jaw, EL = eye length, SE = snout–eye, HL = head length, Dorsal = dorsal fin length, Caudal = caudal fin length, PD = caudal peduncle depth, PL = caudal peduncle length, Pelvic = pelvic fin length, Anal = anal fin length, Pectoral = pectoral fin length. Principal components, PC1 and PC2, of Atlantic salmon explained 24.16% and 14.75% of the variance, respectively, whereas PC1 and PC2 of sea trout explained 25.04% and 22.82% of the variance. Groups identified on the PCAs are represented as follows: • = successful migrants, × = non-successful migrants.
